# A microsporidial deubiquitinase blocks ubiquitin transfer from adenylated E1 to human UBE2K ubiquitin conjugating enzyme

**DOI:** 10.1101/2025.07.31.666823

**Authors:** Tomiwa Lawal, Alana H. Chang, Lauren G. Carnley, Nehry Y. Celoge, Taylor A. Blount, Cynthia Vied, Antonia A. Nemec, Robert J. Tomko

## Abstract

Intracellular pathogens frequently subjugate the ubiquitin system to evade host immune defenses and establish intracellular replication niches. Microsporidia are obligate intracellular animal parasites that typically cause self-limiting infections in humans, but can sometimes cause fatal disseminated disease. At present, the ubiquitin system of microsporidia is virtually unexplored. Here, we discover a likely effector deubiquitinase (DUB) of the otubain subgroup from the human pathogenic microsporidian *Encephalitozoon hellem*, which we designate ehOTUB1. We find that ehOTUB1 selectively binds the human ubiquitin conjugating (E2) enzyme UBE2K and inhibits its ubiquitin conjugation activity independent of ehOTUB1 DUB activity. We show that ehOTUB1 obstructs docking of UBE2K onto ubiquitin E1 enzyme via steric conflict with ubiquitin in the E1 adenylation site to prevent ubiquitin transfer to UBE2K. This unconventional mechanism of E2 inhibition expands the known repertoire by which pathogens manipulate ubiquitin signaling, and suggests that direct inhibition of E2 enzymes may be a broader function of otubain subfamily DUBs than originally appreciated.

## Introduction

Microsporidia are eukaryotic obligate intracellular parasites that can infect most animal phyla, including that of humans^1,2^. They are distantly related to Fungi, and typically spread as infectious spores that are transmitted by ingestion of contaminated food or water. These spores can remain infectious for years under ideal conditions. Microsporidial infections have significant agricultural, aquacultural, and industrial impacts^3^ and are of increasing concern for human health^1^. It is estimated that approximately 10.5% of humans are infected with microsporidia^4^. Infection typically occurs via the gastrointestinal tract, and causes self-limiting diarrhea or is asymptomatic in most immunocompetent patients^1^. However, in the immunocompromised and in some immunocompetent patients, infections can become disseminated and can be fatal^5–7^. Despite their economic and health impacts, very little is known about microsporidial biology and no broadly effective drugs exist to treat microsporidiosis.

Microsporidian species of the genus *Encephalitozoon* are one of the most common causes of infection in humans^1^. Like all microsporidia, *Encephalitozoon* species possess a unique infection organelle called a polar tube that penetrates the host cell membrane and injects the infectious sporoplasm into the host cell cytoplasm^8^. There, the sporoplasm replicates as meronts and sporonts that mature into infectious spores. The precise course of parasite development and replication inside the host differs among species of microsporidia, but normally culminates in the rupture of the host cell and the release of infectious spores into the environment to infect additional hosts or cells^1^.

A common feature of intracellular pathogens is the delivery of effector proteins into the host cell to facilitate pathogen immune evasion and establishment of an intracellular replication niche. In many cases, such effectors modulate the function of the ubiquitin-proteasome system to accomplish these goals^9–11^. The ubiquitin-proteasome system controls diverse cellular processes via the regulated attachment and removal of a small protein, ubiquitin, from other biomolecules (usually proteins)^12^. Typically, ubiquitin is attached to lysine epsilon amino groups or the alpha amino group on proteins via the sequential actions of E1, E2, and E3 enzymes. The E1 enzyme activates the C-terminus of ubiquitin for conjugation via an initial adenylation reaction that ultimately generates a ubiquitin thioester on the E1 catalytic cysteine. An E2 enzyme then associates with the thioesterified E1, prompting ubiquitin transfer to the E2 catalytic cysteine. In a final step, ubiquitin is transferred onto a protein substrate, in a process frequently mediated by an E3 enzyme. The E3 enzyme brings the E2 and substrate in close proximity and catalyzes ubiquitin transfer. The initial ubiquitin can then be modified by additional ubiquitin molecules to form a polyubiquitin chain.

Ubiquitin’s seven lysines (K6, K11, K27, K29, K33, K48, K63) and alpha amino group (M1) permit eight linkage types^13^. Each linkage adopts distinct three-dimensional conformations, permitting the recruitment of linkage-selective ubiquitin-binding proteins that couple a given linkage to specific downstream processes. For example, K63-linked polyubiquitin regulates endocytosis and DNA damage signaling^14^, whereas K48-linked polyubiquitin typically drives protein degradation by the proteasome, and helps activate the non-canonical NFκB pathway^15^.

The attachment of ubiquitin to proteins can be reversed by the actions of deubiquitinating enzymes (DUBs). To date, eight families of DUBs have been identified^16–19^, with the majority being CA-clan cysteine proteases harboring a papain fold^20^. Importantly, many viral and bacterial pathogens encode DUBs^21–26^; these proteases typically suppress ubiquitin-dependent NFκB or IRF signaling by the host cell or reprogram the intracellular environment to favor pathogen invasion, replication, and dissemination. Whereas much has been learned about viral and bacterial enzymes that modulate ubiquitin signaling, very little is known about the ubiquitin-proteasome system of microsporidia or how they may modulate host cell ubiquitination during the infectious cycle^27–29^, especially in mammalian host cells^30^.

To address this knowledge gap, we performed phylogenetic and proteomic searches for microsporidial DUBs encoded by the human pathogenic microsporidian *Encephalitozoon hellem*. We identified a reduced complement of DUBs compared to that found in other simple eukaryotes. We investigated one such DUB here named ehOTUB1, which lacks an obvious ortholog in many lineages of microsporidia as well as many eukaryotes. We show that ehOTUB1 has a strong preference for K48 linkages and is indiscriminate of length or other topologies. Unexpectedly, we find that ehOTUB1 specifically associates with human E2 enzyme UBE2K and blocks its ubiquitin chain conjugation activity in a manner that is independent of ehOTUB1 DUB activity. Using a combination of molecular modeling, mutagenesis, and site-specific crosslinking, we show that ehOTUB1 preferentially blocks association of UBE2K with ubiquitin E1 enzyme harboring ubiquitin at the E1 adenylation site, thereby preventing ubiquitin transfer from transthiolation-competent E1 to UBE2K. Our data thus describe an unusual mechanism for counteraction of E2 activity by a parasite DUB, and provide a mechanistic framework for investigating ehOTUB1 as a drug target for treatment of *Encephalitozoon* infections.

## Results

### Survey of Encephalitozoon hellem deubiquitinating enzymes reveals a conserved otubain DUB

We performed a phylogenetic search of the *E. hellem* genome^31^ for members of the eight families of DUBs. We identified: i) one JAMM metalloproteinase with high sequence identity to the proteasome subunit Rpn11; ii) five members of the ubiquitin-specific protease family; and iii) one member of the ovarian tumor (OTU) family (Table 1). We also identified a Ulp family member that likely cleaves the ubiquitin-like protein <u>s</u>mall <u>u</u>biquitin-related <u>mo</u>difier (SUMO), and a putative protease related to the protein DeSI, which has been reported to cleave SUMO^32^. Of the five USP family members, we could confidently assign them based on sequence homology as orthologs of yeast *DOA4*, *UBP8, UBP12, UBP14,* and *UBP15*. However, we could not confidently assign the OTU family member EHEL_050640 as an obvious ortholog of a *S. cerevisiae* or human DUB based on sequence homology or on domain architecture. EHEL_050640 shared weak sequence homology to human OTUB1 and OTUB2, suggesting it was a member of the otubain subfamily of OTU DUBs^33^.

We next attempted to validate our phylogenetic search using a proteomic approach. Ubiquitin C-terminal propargylamides (Ub^prg^) serve as suicide inhibitors of cysteine DUBs, reacting with the active site thiolate to yield a covalent, stable vinyl thioether^34,35^. The resultant thioalkyl bond is highly stable, permitting purification of any DUBs that have reacted with Ub^prg^ under stringent conditions. We immobilized Halo-tagged Ub^prg^ on agarose beads, and reacted it with lysates from 293T cells that were either infected or uninfected with *E. hellem* (Fig. 1A). After extensive washing, Ub^prg^-protein conjugates were eluted using human rhinovirus 3C protease and analyzed by LC-MS/MS.

**Figure 1:**
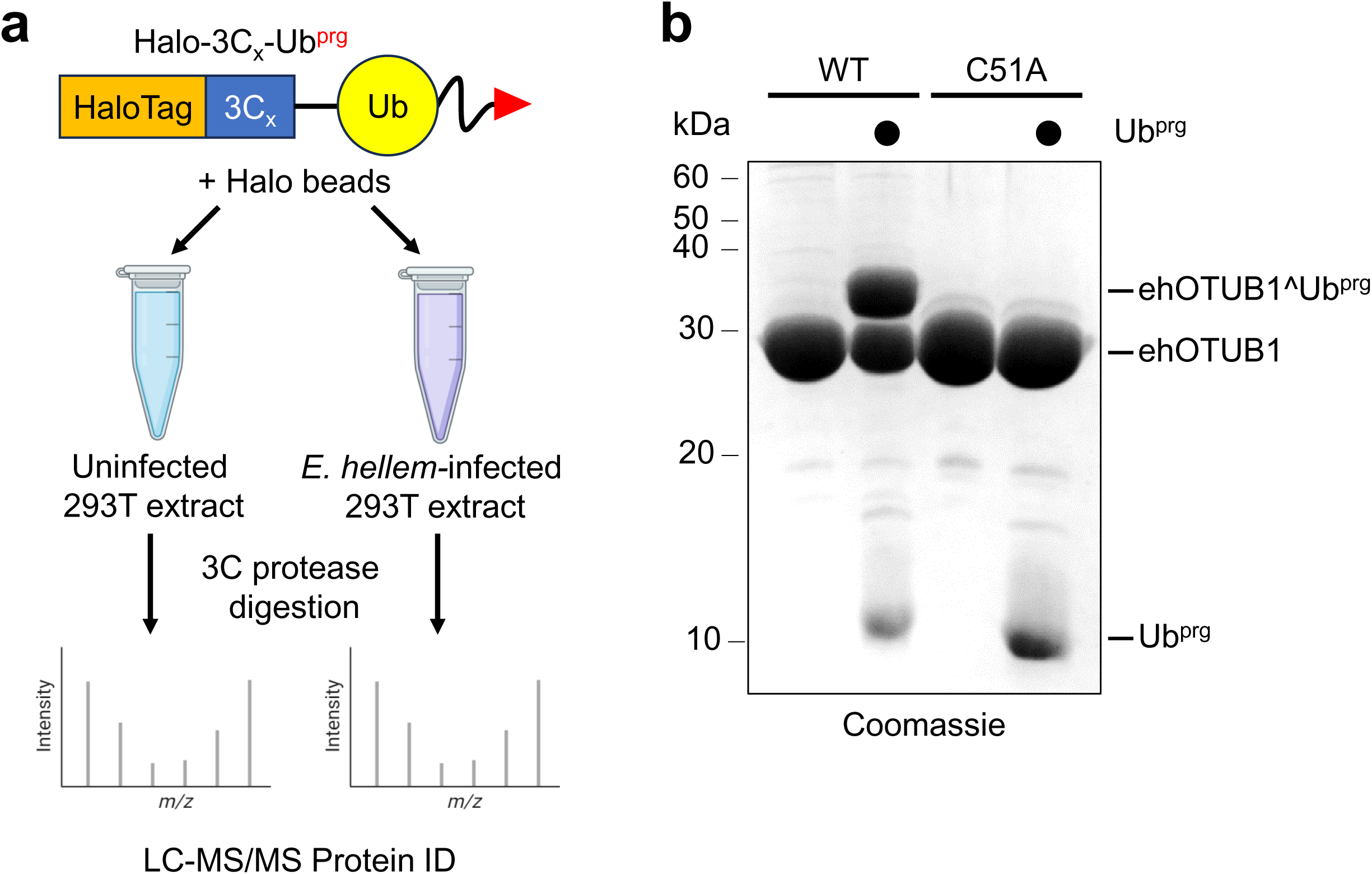
Identification of EHEL_050640 as a putative microsporidial deubiquitinating enzyme. ***A***, Scheme for covalent trapping and mass spectrometric identification of microsporidial deubiquitinating enzymes. A ubiquitin propargylamide suicide probe immobilized via an N-terminal Halo tag was incubated with extracts of uninfected or *E. hellem* infected 293T cells, and after extensive washing, ubiquitin and any covalently bound proteins were eluted using human rhinovirus 3C protease for identification via LC-MS/MS. ***B***, Purified recombinant WT ehOTUB1, but not ehOTUB1(C51A), reacted with ubiquitin propargylamide (Ub^prg^).

As anticipated, several human DUBs were identified in both samples; however, only the infected sample contained *E. hellem* DUBs (Table 2). From this list, we decided to investigate EHEL_050640 for the following reasons: i) it had no obvious direct ortholog in humans or *S. cerevisiae*; ii) a previous report localized this protein to the microsporidial outer membrane and cell wall where it is positioned to modify ubiquitinated proteins within the host cell^36^; and iii) OTU family DUBs are common effector proteins produced by pathogenic human viruses and bacteria^22,23,25,26^. Because EHEL_050640 showed highest sequence homology to the otubain (OTUB) subfamily of OTU DUBs vs. other OTU family members, we hereafter refer to this gene product as ehOTUB1.

### Substrate profiling of ehOTUB1 reveals preferences for K48-linked ubiquitin chains

We first confirmed that ehOTUB1 harbored DUB activity. We expressed and purified ehOTUB1 from *E. coli*, along with a mutant ehOTUB1 in which the predicted catalytic cysteine was substituted with alanine (C51A) (Supplementary Fig. S1A). Covalent reaction with Ub^prg^, evident as a high-molecular weight adduct by SDS-PAGE, was observed for ehOTUB1, but not ehOTUB1(C51A) (Fig. 1B), suggesting ehOTUB1 was a *bona fide* DUB that required Cys51 for catalysis. In further support, ehOTUB1 could cleave the fluorogenic substrate Ub-AMC, and ehOTUB1 activity was reduced by preincubation with the DUB inhibitor ubiquitin aldehyde (Supplementary Fig. S1B).

Some DUBs can process both ubiquitin and other ubiquitin-like proteins (Ubls); for example, the papain-like proteases of coronaviridae display robust activity against both ubiquitin and the interferon-stimulated Ubl ISG15, as well as less potent activity against another Ubl, NEDD8^26,37^. We thus used a series of Ubl^prg^ probes to characterize the selectivity of ehOTUB1 for different Ubls. As shown in Figure 2A (red arrows), ehOTUB1 reacted readily with Ub^prg^ and NEDD8^prg^, but only very weakly with an ISG15-derived probe. No reactivity was observed with SUMO-1^prg^ or SUMO-3^prg^ under any conditions tested. We confirmed the selectivity of ehOTUB1 using fluorogenic probes for Ub-, NEDD8-, and ISG15-hydrolase activity. Cleavage of Ub-AMC was detectable upon incubation with as little as 100 nM ehOTUB1 (Fig. 2B), whereas ehOTUB1 cleaved NEDD8-Rhodamine only at high enzyme concentrations (Fig. 2C), and did not cleave ISG15-AMC even at 10 μM ehOTUB1 (Fig. 2D). Thus, ehOTUB1 prefers ubiquitin over other Ubls.

**Figure 2:**
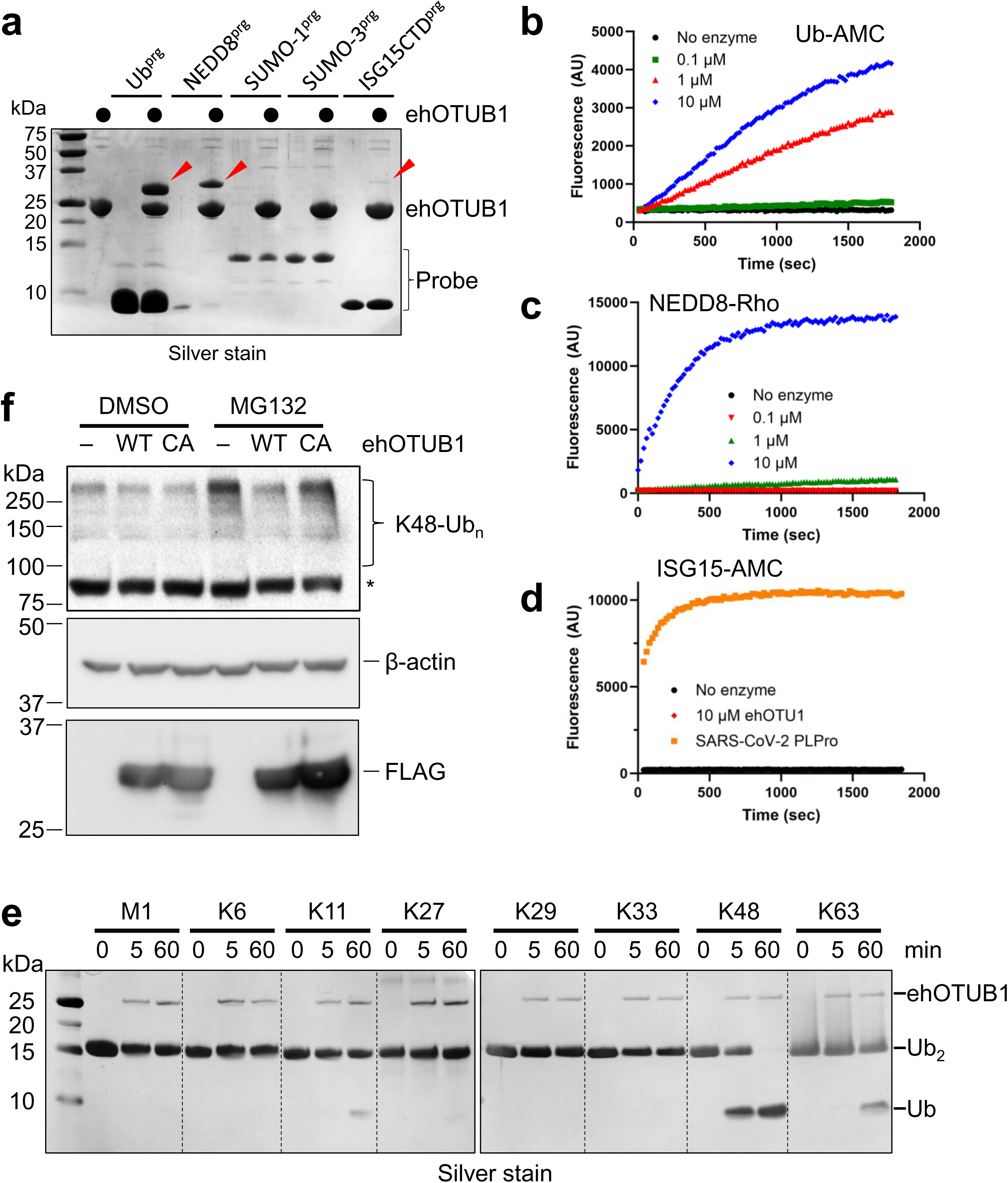
Characterization of EHEL_050640/ehOTUB1. ***A***, ehOTUB1 was reacted with propargylamides of the indicated ubiquitin-like proteins to identify possible substrates. Red arrowheads indicate ehOTUB1 covalently modified with the ubiquitin-like protein probes. ***B-D***, The indicated concentrations of ehOTUB1 were incubated with fluorogenic substrates ubiquitin-AMC (Ub-AMC) (***B***), NEDD8-rhodamine 110 (***C***), or ISG15-AMC (***D***) and fluorescence liberated was measured over time. SARS-CoV-2 papain-like protease (PLPro; 100 nM) was included as a positive control for ISG15-AMC hydrolysis. ***E***, 500 nM ehOTUB1 was incubated with 20 µM Ub_2_ harboring the indicated linkages for 0, 5, or 60 minutes before quenching of the reactions and analysis by SDS-PAGE. ***F***, Flp-In T-REx 293 cells were transfected with plasmids encoding the indicated forms of ehOTUB1, and after 24 hours, were treated with 20 µM MG132 or DMSO control for 4 hours prior to analysis. An asterisk indicates an apparent nonspecific band.

To determine whether ehOTUB1 displayed ubiquitin linkage preferences, we assayed its ability to cleave diubiquitin (Ub_2_) harboring each of the eight possible linkages. As shown in Fig. 2E, ehOTUB1 preferentially cleaved K48-linked Ub_2_, with greater than 50% of the substrate cleaved within 5 minutes. Upon prolonged incubation, some weak cleavage of K11- and K63-linked Ub_2_ was evident. No changes in linkage selectivity were observed against longer chains (Supplementary Fig. S2A), and ehOTUB1 DUB activity was not appreciably altered against branched Ub_3_ chains (Supplementary Fig. S2B), suggesting ehOTUB1 primarily recognizes one or both ubiquitin moieties flanking the scissile K48 isopeptide bond rather than a particular chain length or other, more complex chain architectures.

We next evaluated whether expression of ehOTUB1 led to cleavage of pUb conjugates in cells. K48-linked ubiquitin chains are lowly abundant in HEK293 cells due to their processing by the proteasome, but are stabilized by a brief treatment of cells with the proteasome inhibitor MG132. We thus treated HEK293 cells expressing empty vector, 3xFLAG-tagged ehOTUB1, or 3xFLAG-ehOTUB1(C51A) with DMSO control or MG132, followed by immunoblotting with K48-specific antibodies. As shown in Fig. 2F, MG132 caused an accumulation of K48-linked ubiquitin chains in the empty vector-expressing cells, and this accumulation was near-completely lost in cells expressing 3xFLAG-ehOTUB1. In contrast, the abundance of K48-linked chains was only slightly decreased in cells expressing 3xFLAG-ehOTUB1(C51A), indicating that ehOTUB1 is catalytically active when produced in human cells, and decreases K48-linked polyubiquitin in a manner primarily dependent upon its catalytic activity. Thus, ehOTUB1 functions as a K48-selective DUB both *in vitro* and in cells.

### Selective inhibition of human UBE2K ubiquitin ligase activity by ehOTUB1

At present, it is not possible to genetically manipulate microsporidia to investigate gene function^38^. However, while searching for orthologs of ehOTUB1, we tested its ability to complement loss of OTU proteins from budding yeast. Of the two *S. cerevisiae* OTU DUBs *OTU1* and *OTU2*, ehOTUB1 had higher sequence similarity to the catalytic domain of the Cdc48-associated DUB *OTU1*, but clearly lacked the N-terminal ubiquitin-like domain conserved across fungal *OTU1* orthologs that mediates interaction with Cdc48^39^. Consistent with this, we failed to copurify Cdc48 with ehOTUB1 when it was expressed in *S. cerevisiae* (Supplementary Fig. S3), and we did not detect Cdc48 or any Cdc48 pathway components in ehOTUB1 pulldowns by mass spectrometry (discussed below). However, expression of N-terminally 3xFLAG-tagged ehOTUB1 even in wild-type yeast caused a substantial growth defect (Fig. 3A) not observed when yeast Otu1 or Otu2 were overproduced. To our surprise, growth suppression of equal magnitude was also observed upon expression of the C51A mutant, indicating that the growth defect was independent of ehOTUB1 DUB activity.

**Figure 3:**
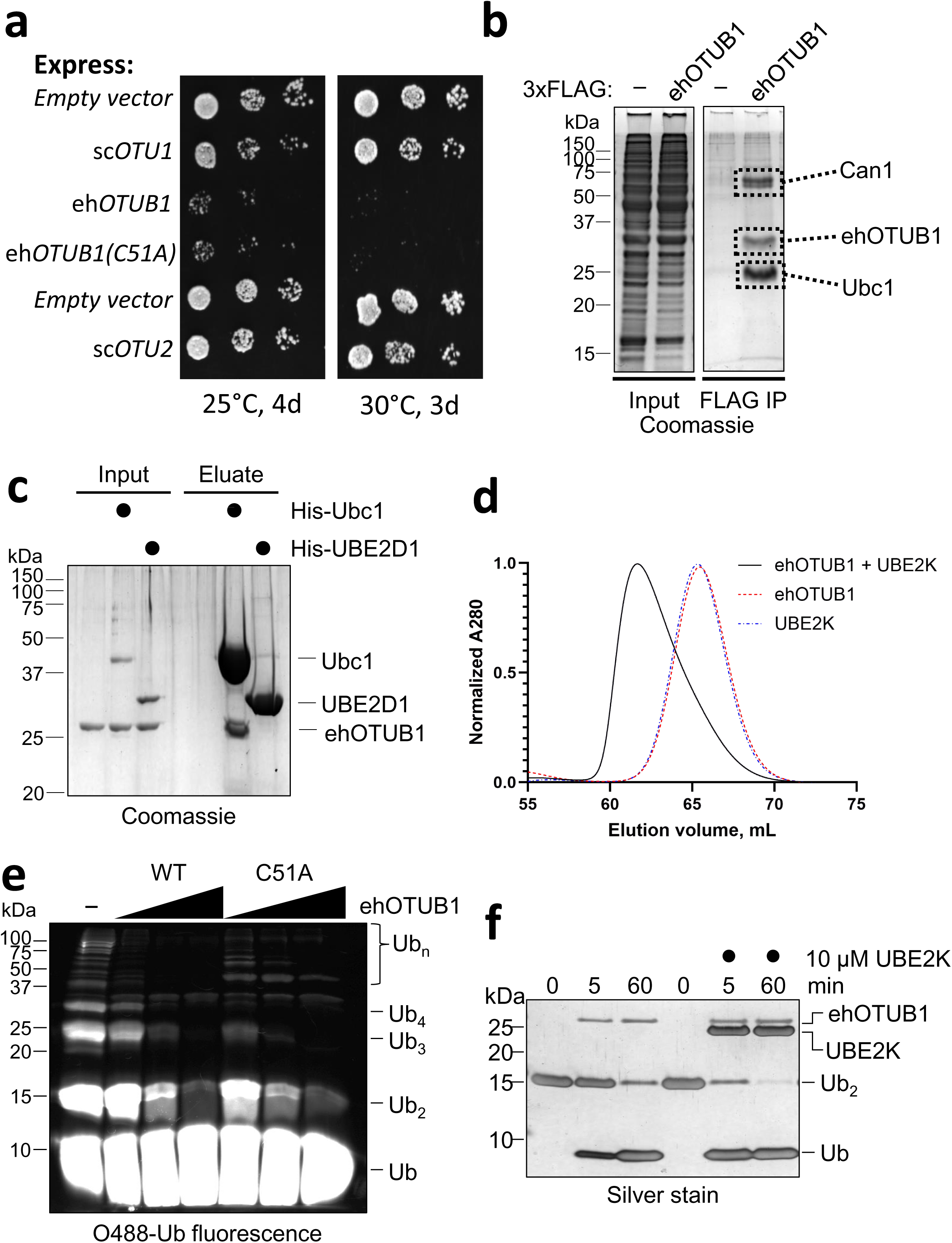
Identification of Ubc1/UBE2K as a target of ehOTUB1-mediated inhibition. ***A***, Cells expressing the indicated proteins as N-terminal 3xFLAG fusions from high-copy vector p424GPD were spotted in six-fold serial dilutions before incubation as shown. ***B***, ehOTUB1 copurifies from yeast with Ubc1. 3xFLAG-ehOTUB1 and interacting proteins were immunopurified from wild type yeast by 3xFLAG affinity, and the boxed areas were sent for protein identification by liquid chromatography-mass spectrometry. The most abundant protein in each band is shown. The input and FLAG IP are non-adjacent lanes from the same gel. ***C***, Ubc1 interacts with ehOTUB1 *in vitro*. Untagged ehOTUB1 was incubated with buffer, His-tagged Ubc1, or His-tagged human UBE2D1 before Ni-NTA affinity purification and SDS-PAGE. ***D,*** Human UBE2K binds ehOTUB1. Equal amounts of ehOTUB1, UBE2K, or ehOTUB1-UBE2K mixture were separated by gel filtration. The leftward shift in the elution volume for the mixture indicates direct interaction. ***E***, ehOTUB1 inhibits formation of K48-linked polyubiquitin chains by UBE2K in a manner independent of ehOTUB1 catalytic activity. Ubiquitination reactions containing 1 µM E1, 5 µM UBE2K, and fluorescent ubiquitin were incubated for two hours in the presence or absence of 1 to 4-fold molar excess of ehOTUB1 or ehOTUB1(C51A) to UBE2K, followed by analysis by SDS-PAGE and fluorescence imaging. ***F***, UBE2K does not interfere with ehOTUB1 deubiquitination activity. 500 nM ehOTUB1 was preincubated with 10 µM human UBE2K prior to addition of 20 µM K48-Ub_2_ for the indicated times.

To provide insight into its biological functions and the cause of its toxicity, we immunopurified ehOTUB1 from wild type (WT) yeast cells using anti-FLAG affinity, and performed gentle elution of bound proteins with 3xFLAG competitor peptide. Analysis of the eluted proteins by SDS-PAGE revealed three prominent species uniquely present in ehOTUB1 pulldowns. LC-MS/MS identified the band migrating at ∼30 kDa as 3xFLAG-tagged ehOTUB1. A second set of closely migrating bands at ∼62 kDa was enriched for the arginine permease Can1 compared to the control sample, and a final species of ∼25 kDa was enriched for the ubiquitin-conjugating enzyme Ubc1, the yeast ortholog of human UBE2K. Ubc1/UBE2K is unique among ubiquitin E2 enzymes in that it is the only E2sthat harbors a C-terminal ubiquitin-associated (UBA) domain; interestingly, we were unable to identify either a Ubc1/UBE2K ortholog or a UBA domain-containing E2 enzyme in any species of microsporidia with a sequenced genome, suggesting that ehOTUB1 is unlikely to bind a microsporidian ortholog of Ubc1/UBE2K.

We next confirmed the interaction between Ubc1 and ehOTUB1 using recombinant purified proteins. As shown in Fig. 3C, untagged ehOTUB1 readily copurified with His-tagged Ubc1, but not with a second E2 enzyme, His-tagged human UBE2D1. We also observed interaction of ehOTUB1 directly with human UBE2K, as evidenced by formation of a faster-eluting complex by size exclusion chromatography (Fig. 3D). As *E. hellem* typically infects mammals^40^, we next investigated the impact of ehOTUB1 on the activity of UBE2K. Like Ubc1, UBE2K exclusively synthesizes K48-linked ubiquitin chains^41^, which matches the cleavage preference of ehOTUB1. We envisioned that ehOTUB1 may thus counteract K48 ubiquitination by UBE2K via chain cleavage. To test this possibility, we added increasing amounts of ehOTUB1 or catalytically inactive ehOTUB1(C51A) to a ubiquitination reaction containing fluorescent ubiquitin, E1 enzyme, and UBE2K (Fig. 3E). In the absence of ehOTUB1, UBE2K synthesized high molecular weight ubiquitin chains (Ub_n_) as well as Ub_2_, Ub_3_, and Ub_4_. However, as ehOTUB1 was titrated in at increasing molar ratios to UBE2K, formation of all ubiquitin chain lengths was reduced in a concentration-dependent manner. Importantly, and consistent with the observation that catalytically inactive ehOTUB1 remained toxic to yeast cells, titration of ehOTUB1(C51A) also led to a concentration-dependent suppression of polyubiquitin formation. Interestingly, some ubiquitin species appeared to be more heavily suppressed by WT ehOTUB1 than the C51A mutant. This suggests that, whereas ehOTUB1 may suppress K48 chain formation in part via its deubiquitinating activity, it also inhibits UBE2K via a non-catalytic mechanism, likely via its direct interaction with UBE2K. Conversely, addition of an ∼20-fold molar excess of UBE2K did not block ehOTUB1 activity; in fact, a small but reproducible enhancement of activity was observed (Fig. 3F), similar to that observed for human OTUB1^42^. Together, these data indicate that whereas ehOTUB1 may suppress K48 chain formation in part via its deubiquitinating activity, it also inhibits UBE2K via a non-catalytic mechanism.

### The mechanism of ehOTUB1 substrate recognition and UBE2K inhibition differs from that of human OTUB1

To our knowledge, a single example of direct E2 inhibition by a DUB (e.g., independent of DUB activity) exists^43^. However, the growth suppression by ehOTUB1(C51A) and inhibition of E2-dependent polyubiquitin chain synthesis was reminiscent of a phenotype observed when human OTUB1 was expressed in yeast^44^. This phenotype was attributed to inhibition of multiple E2 enzymes, including orthologs of UBE2N−UBE2V1 (Ubc13-Mms2) and UBE2D family members^44–46^, but no report investigating human OTUB1 has linked it to regulation of UBE2K. Notably, inhibition of UBE2N-UBE2V1 by human OTUB1 was potentiated by inclusion of ubiquitin aldehyde^45^, which occupied the distal ubiquitin (e.g., the ubiquitin whose C-terminus is conjugated to an amino group; see cartoon in Fig. 4E) binding site of human OTUB1 to mimic its interaction with a native K48-ubiquitin linkage. However, ehOTUB1 had no effect on the formation of K63 chains by UBE2N−UBE2V1 in the presence or absence of ubiquitin aldehyde (Supplementary Fig. S4), further supporting an apparently distinct E2 selectivity of ehOTUB1 vs. human OTUB1.

**Figure 4:**
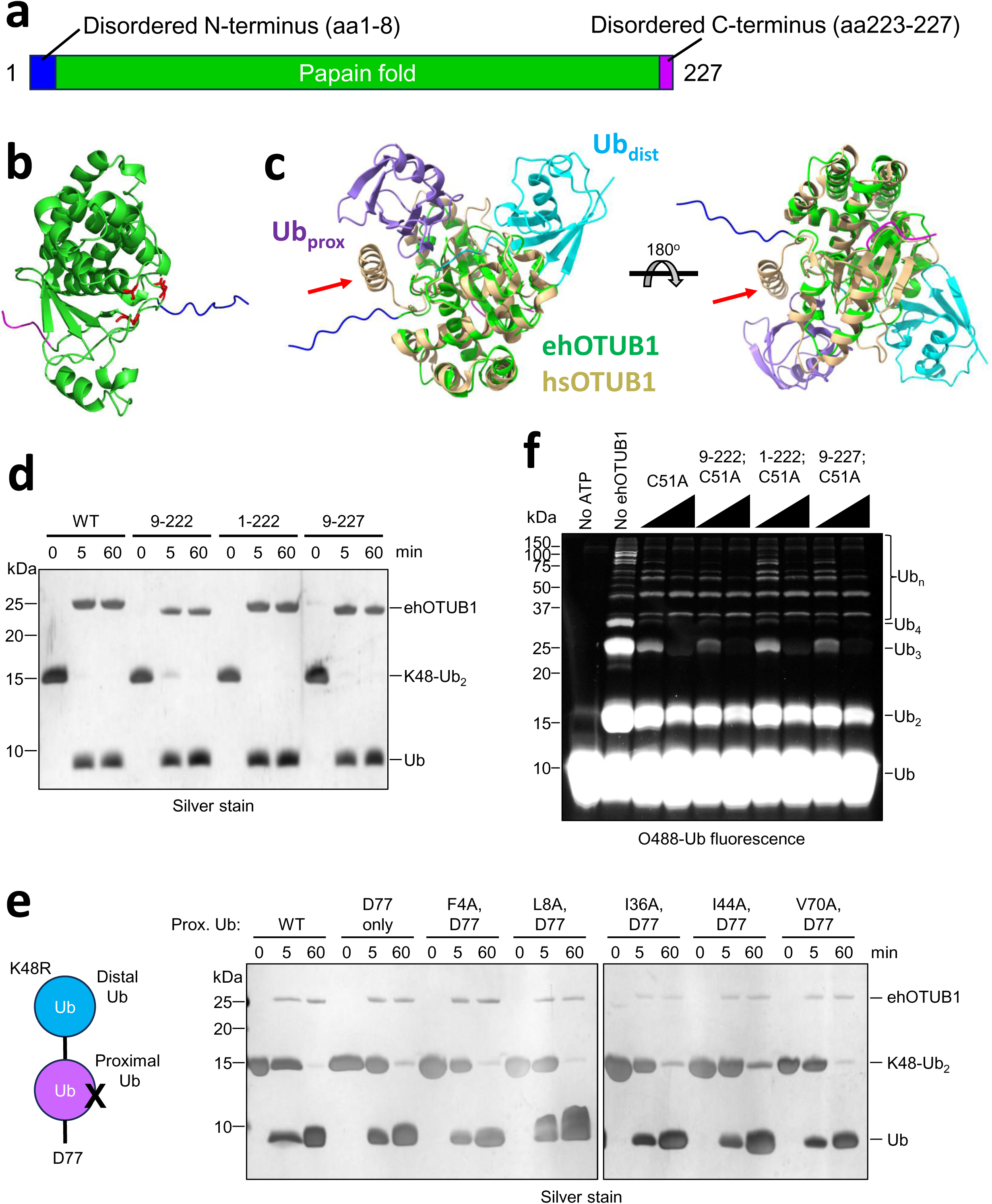
The N- and C-termini of ehOTUB1 are dispensable for ehOTUB1 functions. ***A***, Domain architecture of ehOTUB1. ***B***, An AlphaFold prediction of the ehOTUB1 structure indicates the extreme N- and C-termini (blue and magenta, respectively) are likely to be disordered. The residues comprising the catalytic triad are shown in red stick mode. ***C***, Superimposition of ehOTUB1 colored as in (A) with human OTUB1 (tan) bound to two ubiquitin molecules (cyan and purple). The human OTUB1 N-terminal helix that binds the proximal ubiquitin is indicated with a red arrow. ***D,*** N- and C-terminal truncation mutants of ehOTUB1 cleave K48-Ub_2_ as well as WT ehOTUB1. K48-Ub_2_ (20 µM) was incubated with 1 µM of the indicated ehOTUB1 proteins at 37°C for the times shown. ***E,*** ehOTUB1 DUB activity is unaffected by mutation of common contact patches on the proximal Ub. As in (***D***), but 500 nM ehOTUB1 was used. ***F***, N- and C-terminal truncation mutants of ehOTUB1 inhibit UBE2K as well as WT. Reactions were performed as in Fig. 3E.

Human OTUB1 requires a conserved N-terminal helix both for its DUB activity and for E2 inhibition^44,45,47^. This helix orients the proximal ubiquitin (e.g., the ubiquitin donating the amino group to the isopeptide bond) during deubiquitination and also confers preference for inhibiting thioesterified UBE2N over *apo* UBE2N via interaction with the thioesterified ubiquitin attached to UBE2N. Recognition of the proximal and thioesterified ubiquitin is via hydrophobic interactions involving ubiquitin L8, I44, and V70 among others. In contrast, ehOTUB1 appears to lack such an N-terminal helix, but is predicted to possess short N- and C-terminal disordered regions (amino acids 1-8 and 223-227, respectively; Fig. 4A) adorning the OTU papain fold that may serve an analogous purpose. In support of this, superposition of an ehOTUB1 Alphafold model (Fig. 4B) onto a structure of ubiquitin aldehyde–OTUB1:UbcH5b–ubiquitin^44^ (where − indicates a covalent bond and : indicates a noncovalent interaction) positioned this N-terminal extension closely to the human OTUB1 helix (Fig. 4C, red arrows).

We thus generated N-, C-, and doubly N/C-terminally truncated ehOTUB1 mutants to assess whether these termini contributed to deubiquitination or to inhibition of UBE2K. All three truncation mutants cleaved K48-Ub_2_ as efficiently as full-length ehOTUB1, indicating these termini were dispensable for substrate positioning and for catalysis (Fig. 4D). In further support, we engineered K48-Ub_2_ mutants in which the proximal ubiquitin (Fig. 4E, purple) harbored mutations in residues that directly contact the human OTUB1 N-terminal helix (e.g., L8A, I44A, V70A)^44,45^. This was accomplished by introduction of K48R into the distal ubiquitin and inclusion of a C-terminal aspartate (D77) onto the proximal ubiquitin to ensure unidirectional conjugation as performed previously^48^. However, ehOTUB1 cleaved each of these Ub_2_ mutants as efficiently as WT K48-linked Ub_2_. Cleavage was also unaffected by mutations disrupting other surfaces commonly mediating ubiquitin recognition (e.g., F4A, I36A; Fig. 4E)^13^; thus, if the proximal ubiquitin is recognized by ehOTUB1, it is likely via an alternative surface. Truncation of the N- and C-termini of ehOTUB1 also had no impact on inhibition of UBE2K (Fig. 4F). Thus, the mode of substrate recognition and E2 inhibition is fundamentally different than that observed previously for human OTUB1.

### AlphaFold modeling identifies hydrophobic interactions mediating ehOTUB1–UBE2K binding

We used AlphaFold3 to predict the structure of the ehOTUB1**–**UBE2K complex (Fig. 5A) to better understand the potential mode of UBE2K inhibition. The five top models could be near-perfectly superimposed, with RMSD < 0.4 Å (Supplementary Fig. S5A), and UBE2K within these structures could be superimposed with RMSD < 0.3 Å onto a published structure of UBE2K^49^. In the ehOTUB1-UBE2K model, a hydrophobic loop of UBE2K docks into a similarly hydrophobic cleft formed between helices 1, 4, and a loop extending from helix 6 of ehOTUB1 (Fig. 5B). The contacts between these two regions are primarily via aromatic and aliphatic side chains, with the UBE2K Phe68 side chain fitting knob-in-hole into a recess formed by ehOTUB1 F99, Y123, and F167. This interaction places the UBE2K A102 and A103 side chains flush against the surface of ehOTUB1, with ehOTUB1 L94 docking into a more distal recess in UBE2K where it interacts with the aliphatic portion of the UBE2K R11 side chain.

**Figure 5:**
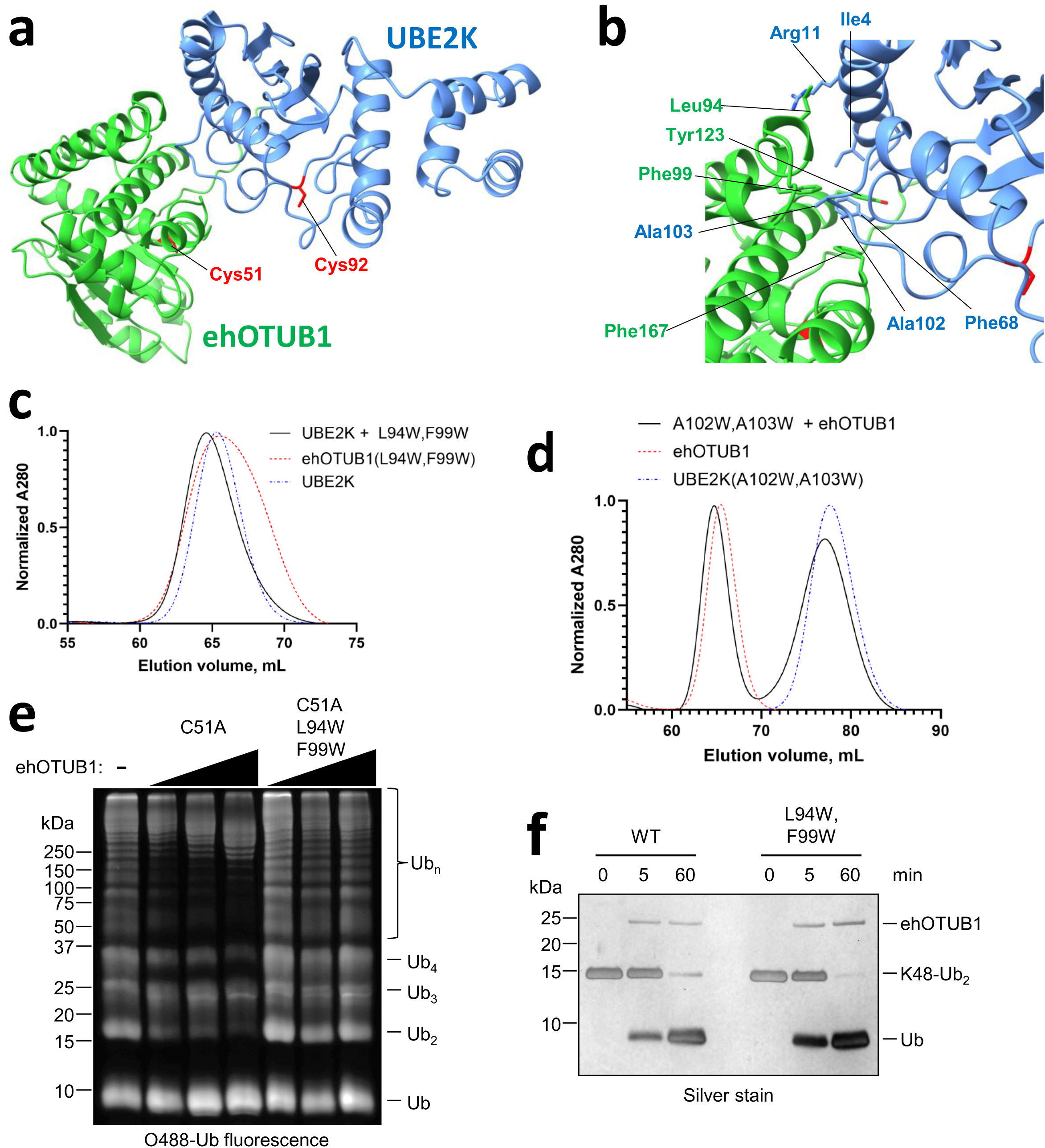
A mutagenesis-validated AlphaFold model of the ehOTUB1−UBE2K complex. ***A***, AlphaFold model of ehOTUB1 (green) in complex with UBE2K (cornflower blue) indicates that the catalytic cysteines (red sticks) of both enzymes remain exposed. ***B***, Detail of the predicted hydrophobic interface between ehOTUB1 and UBE2K. Residues at the interface between the two proteins are indicated. Catalytic cysteines are shown as red sticks. ***C*** and ***D***, Mutation of residues at the predicted ehOTUB1-UBE2K interface disrupt their interaction. Equal amounts of the indicated WT and mutant proteins or mixtures thereof were separated by gel filtration. Note that introduction of tryptophan residues into UBE2K caused enhanced hydrophobic interactions with the gel filtration resin that slowed its migration relative to the WT protein. ***E***, The ehOTUB1(L94W,F99W) mutant retains WT DUB activity. ***F***, UBE2K ubiquitin conjugation activity is unaffected by the ehOTUB1(C51A,L94W,F99W) mutant.

We validated this mode of interaction via structure-guided mutagenesis. Whereas WT UBE2K co-eluted with WT ehOTUB1 in a faster-migrating complex by size exclusion chromatography (Fig. 3D), it did not do so with ehOTUB1(L94W,F99W), indicating these two proteins did not interact (Fig. 5C, black vs. red and blue traces). UBE2K(A102W,A103W) eluted substantially later than WT UBE2K due to hydrophobic interactions of the tryptophan substitutions with the size exclusion resin (compare Fig. 5C and 5D blue traces); regardless, it failed to induce a migration shift in WT ehOTUB1 (Fig. 5D and see Supplementary Fig. S5B), indicating the two proteins did not interact.

We next used the ehOTUB1(L94W,F99W) UBE2K interaction mutant to assess whether binding to UBE2K was required for UBE2K ubiquitin-conjugating activity. Whereas we again observed concentration-dependent suppression of UBE2K ubiquitin conjugating activity by ehOTUB1(C51A), we observed little to no inhibition of UBE2K by ehOTUB1(C51A,L94W,F99W) (Fig. 5E). The loss of UBE2K inhibition was not due to a general structural defect induced in ehOTUB1 by the L94W or F99W mutations, because ehOTUB1(L94W,F99W) retained WT DUB activity (Fig. 5F). Thus, ehOTUB1 must interact with UBE2K to inhibit its ubiquitin-conjugating activity.

### ehOTUB1 prevents docking of UBE2K onto Ub-loaded E1 to prevent transthiolation of UBE2K

Inspection of the AlphaFold model of the ehOTUB1−UBE2K complex did not reveal an obvious mechanism of inhibition, as the catalytic sites of both enzymes remained fully exposed (Fig. 5A). Consistent with this, superimposition of the ehOTUB1−UBE2K complex onto a structure of UBE2K thioesterified with ubiquitin (Supplementary Fig. S6A)^49^ or poised for chain elongation (Supplementary Fig. S6B)^50^ revealed no obvious steric clash with donor or acceptor ubiquitin. However, we found that ehOTUB1(C51A) prevented the formation of ubiquitin thioester on the catalytic cysteine of UBE2K, evident as a DTT-labile band shift by non-reducing SDS-PAGE (Ub∼UBE2K, Fig. 6A). This inhibition of Ub∼UBE2K formation was dependent on ehOTUB1-UBE2K interaction, because the ehOTUB1(C51A,L94W,F99W) mutant had no effect on the efficiency or kinetics of Ub∼UBE2K formation. We thus focused on the potential for steric clash between E1 and ehOTUB1 as the mechanism of UBE2K suppression by ehOTUB1.

**Figure 6:**
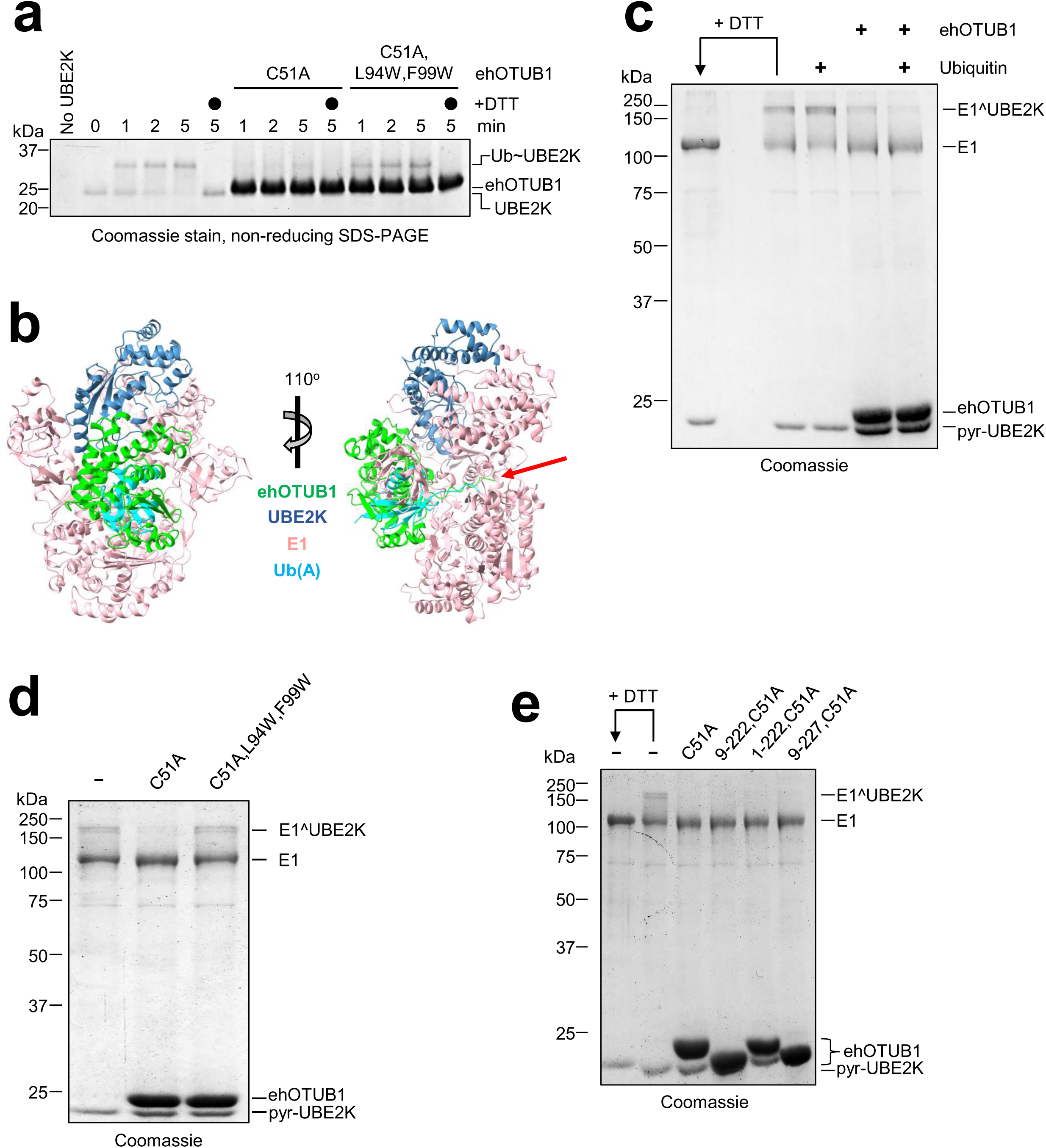
ehOTUB1 interferes with recruitment of UBE2K to ubiquitin-loaded E1 enzyme. ***A***, UBE2K ubiquitin thioester (Ub∼UBE2K) formation is blocked by ehOTUB1(C51A) but not by ehOTUB1(C51A,L94W,F99W). ***B,*** The ehOTUB1−UBE2K AlphaFold model was superimposed onto Ubc4 (omitted for clarity) from PDB 4II2. E1 and the ubiquitin in the adenylation site are shown in pink and cyan, respectively). The potential accessing of the E1 adenylation site by the ehOTUB1 N-terminal region is indicated with a red arrow. ***C***, Inhibition of E1-UBE2K crosslinking by ehOTUB1 is enhanced in the presence of Ub. E1 and disulfide-activated UBE2K were pre-incubated with ubiquitin and ehOTUB1, respectively, where indicated, before mixing and separation by non-reducing SDS-PAGE. The position of the E1-UBE2K disulfide is indicated as E1^UBE2K. ***D,*** The ehOTUB1(C51A,L94W,F99W) mutant fails to suppress the E1-UBE2K crosslink. ***E***, The ehOTUB1 N- and C-termini are dispensable for inhibition of E1-UBE2K crosslinking.

During ubiquitin activation, the E1 activating enzyme first uses ATP and Mg^2+^ to adenylate the C-terminus of ubiquitin at the E1 adenylation site (hereafter, Ub(A) describes ubiquitin bound at the adenylation site). This is followed by rotation of the E1 cysteine half-domain that drives formation of a ubiquitin thioester on the E1 catalytic cysteine (hereafter, Ub(T) describes thioesterified ubiquitin on the E1 catalytic cysteine)^51^. Loading of a new ubiquitin molecule at the adenylation site then stimulates E2 binding and transfer of Ub(T) to the E2 catalytic cysteine for subsequent ubiquitination of a substrate^52^.

At present, no structure of UBE2K in complex with E1 exists; we thus modeled ehOTUB1–UBE2K onto a structure of the closely related E2 enzyme Ubc4 in complex with E1 and Ub(A)^53^ (Fig. 6B). UBE2K superimposed onto Ubc4 with a backbone RMSD of 0.766 Å and no appreciable clash with E1. Similarly, zero steric conflict was observed between ehOTUB1 and E1 aside from the ehOTUB1 flexible N-terminus accessing the E1 adenylation site (Fig. 6B, red arrow). Given that the ehOTUB1 N-terminus was dispensable for UBE2K inhibition (Fig. 4F) it is unlikely to interfere with access of the flexible ubiquitin C-terminus to the adenylation site. In contrast, ehOTUB1 displayed near-complete overlap with the globular domain of Ub(A) in this model (Fig. 6B), suggesting that the UBE2K-ehOTUB1 complex was incompatible for docking onto Ub(A)−E1 complexes. In contrast, analogous modeling of ehOTUB1–UBE2K onto structures of Ubc4 undergoing ubiquitin hand-off from doubly-ubiquitin loaded E1^54^ revealed no clash between ehOTUB1 and Ub(T) (Supplementary Fig. S6C,D), indicating that Ub(A) is likely the major source of steric clash.

To test this, we exploited the close positioning of the E1 and E2 catalytic cysteines necessary for ubiquitin transfer. A disulfide bond can be formed between the E1 and E2 catalytic cysteines if: i) the E2 enzyme is pre-activated for disulfide exchange using the mild oxidant 2,2’-dipyridyldisulfide^53^; and ii) ATP is withheld to prevent Ub(A) transfer to Ub(T), which would in turn block E1-E2 crosslinking. Indeed, addition of 2,2’-dipyridyldisulfide-activated UBE2K (pyr-UBE2K) to E1 under these conditions yielded a band migrating higher than E1 by non-reducing SDS-PAGE, indicative of the E1-E2 crosslink (Fig. 6C). This band was dependent on both pyr-UBE2K and E1 (Supplementary Fig. S7) and was sensitive to DTT as anticipated (Fig. 6C). Inclusion of ubiquitin in the reaction enhanced crosslink formation, presumably by promoting rotation of the E1 cysteine half-domain in response to Ub(A) as reported previously^55,56^.

We next tested the impact of ehOTUB1 on this interaction; we used ehOTUB1(C51A), which could be purified as a well-behaved monomer in the absence of any reducing agent that might otherwise interfere with crosslink formation. Inclusion of ehOTUB1(C51A) in the E1-E2 crosslinking reaction performed in the absence of ubiquitin yielded a reproducible but modest suppression of the crosslink (Fig. 6C). This suggested that some limited steric clash may occur between *apo* E1 and ehOTUB1. Importantly, pre-incubation of E1 with ubiquitin prior to addition of ehOTUB1(C51A)−UBE2K further suppressed the crosslink, despite potentiation of the crosslink by ubiquitin in the absence of ehOTUB1(C51A) (Fig. 6C). This observation agreed closely with our modeling (Fig. 6B), and we thus included ubiquitin in all subsequent crosslinking reactions. Importantly, the interaction mutant ehOTUB1(C51A,L94W,F99W) failed to inhibit crosslink formation even in the presence of excess ubiquitin (Fig. 6D), indicating that interaction between ehOTUB1 and UBE2K was required to inhibit UBE2K docking onto E1. This agreed closely with the failure of ehOTUB1(C51A,L94W,F99W) to suppress UBE2K ubiquitin chain formation (Fig. 5E) or to block Ub∼UBE2K formation (Fig. 6A). As observed for inhibition of UBE2K (Fig. 4F), both the N- and C-termini of ehOTUB1 were dispensable for blocking E1-UBE2K crosslink formation (Fig. 6E).

Although our findings thus far support our model of ehOTUB1 sterically clashing with Ub(A) to inhibit UBE2K, it is known that inhibition of UBE2N−UBE2V1 by human OTUB1 is potentiated by binding of ubiquitin to the human OTUB1 active site^44,45^. To rule out the possibility that ubiquitin docking to the ehOTUB1 active site was potentiating its inhibition as in Fig. 6C, we identified ehOTUB1 residues likely to be critical for docking of ubiquitin into the distal ubiquitin position (Fig. 4C) based on an AlphaFold model of Ub−ehOTUB1 (Fig. 7A). We screened four mutations in ehOTUB1 anticipated to disrupt distal ubiquitin binding, and identified two, Y150A and Y200A, based on complete compromise of K48-Ub_2_ cleavage (Fig. 7B). This compromised cleavage was due to loss of interaction with the distal ubiquitin, because these mutations also compromised cleavage of Ub-AMC (Fig. 7C), and led to loss of comigration of K48-Ub_2_ and ehOTUB1 by gel filtration (Fig. 7D). The complete loss of interaction between these mutants and K48-Ub_2_ supports our prior supposition that ehOTUB1 primarily recognizes the distal ubiquitin (Fig. 4E). Importantly, pre-incubation of pyr-UBE2K with ehOTUB1(C51A,Y150A) or ehOTUB1(C51A,Y200A) impaired both crosslink formation between E1 and UBE2K (Fig. 7E) as well as polyubiquitin chain formation by UBE2K (Fig. 7F) as effectively as ehOTUB1(C51A). Thus, we conclude that ubiquitin binding at the ehOTUB1 active site is not appreciably involved in its inhibition of UBE2K. Instead, we suggest that binding of ehOTUB1 to UBE2K preferentially blocks recruitment of UBE2K to Ub(A)-loaded forms of E1 to prevent transthiolation. This observation is further supported by the lack of detectable E1 enzyme co-purified with 3xFLAG-ehOTU1 in yeast, despite robust copurification of Ubc1 (Fig. 3B).

**Figure 7:**
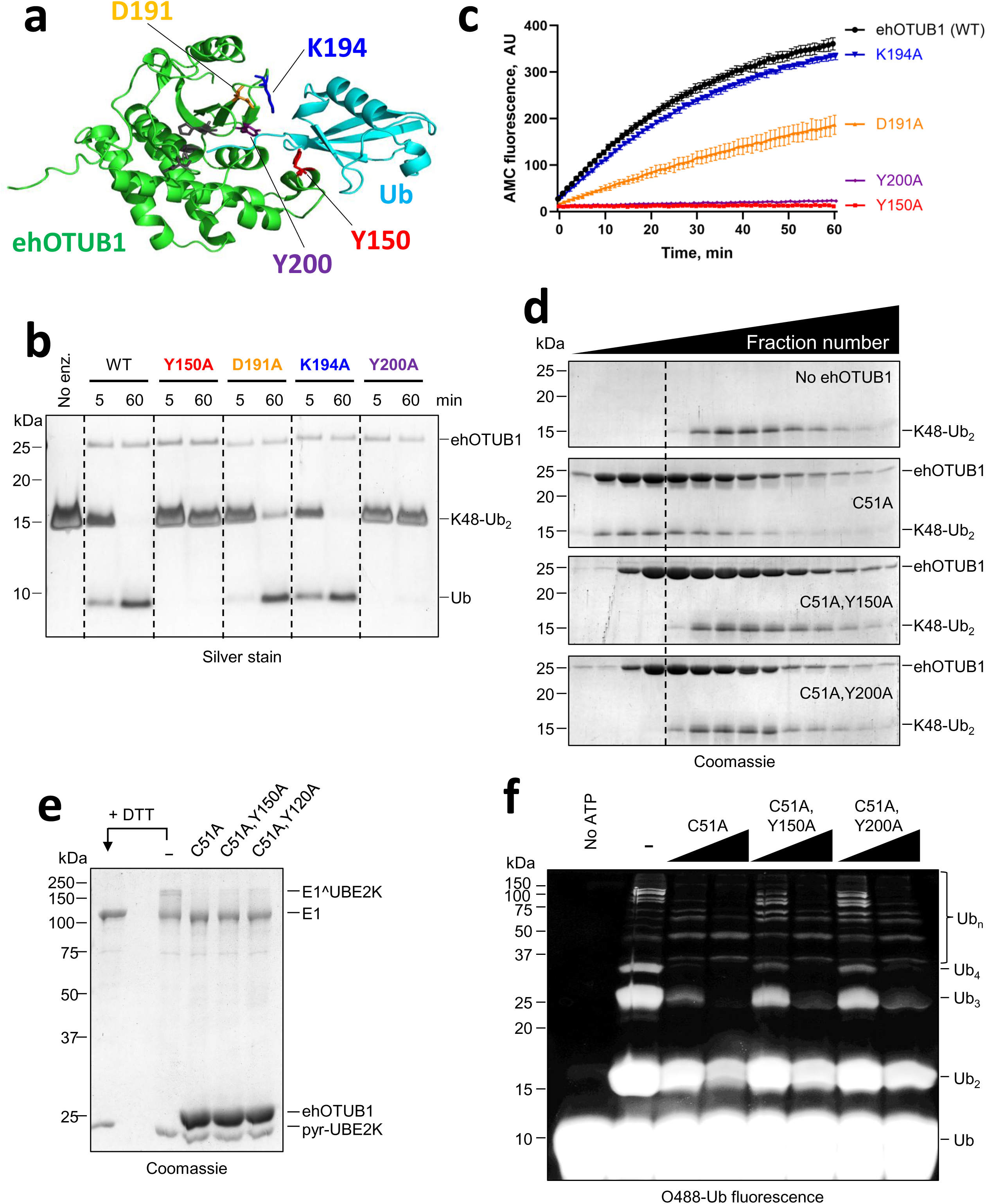
Docking of ubiquitin into the ehOTUB1 active site is dispensable for UBE2K inhibition. ***A***, AlphaFold model of ehOTUB1 (green) in complex with distal ubiquitin (cyan), with residues predicted to contact ubiquitin highlighted. ***B***, The indicated ehOTUB1 proteins (500 nM) were incubated with 20 µM K48-Ub_2_ for the times shown before analysis by SDS-PAGE. ***C***, Ub-AMC hydrolysis assay of WT and mutant ehOTUB1. The indicated ehOTUB1 proteins (500 nM) were incubated with 500 nM Ub-AMC, and fluorescence was monitored over time. ***D***, The Y150A and Y200A ehOTUB1 mutations abrogate interaction with K48-Ub_2_. K48-Ub_2_ (80 µM) was incubated with the indicated ehOTUB1 proteins (90 µM) for one hour at room temperature before separation by gel filtration. ***E***, Y150A and Y200A ehOTUB1 mutants retain the ability to block E1-UBE2K interaction as visualized by crosslinking. ***F***, Y150A and Y200A ehOTUB1 mutants retain the ability to block ubiquitin conjugation by UBE2K.

Taken together, our data suggest that ehOTUB1 sterically clashes with Ub(A)-E1. Given that adenylation of ubiquitin is prerequisite for its thioesterification, and that loading of Ub(A) at the E1 adenylation site is coupled to transthiolation of E2^52,54–56^, we conclude that ehOTUB1 preferentially blocks recruitment of UBE2K to adenylated and presumably also transthiolation-primed E1 to suppress UBE2K signaling.

## Discussion

The importance of ubiquitin signaling to host cell defense is underscored by the numerous intracellular pathogens that encode ubiquitin proteases. Our survey of microsporidial DUBs suggests that *E. hellem* harbors a greatly reduced complement of deubiquitinating enzymes compared even to *S. cerevisiae* and most other single-celled eukaryotes, and has revealed to our knowledge the first known example of a pathogen otubain DUB directly inhibiting a human E2 enzyme. Interestingly, both the E2 preference and the mechanism of E2 inhibition differ significantly from that of human OTUB1, the only other well characterized example of otubain inhibition of an E2^44–46^. However, the observation that ehOTUB1 similarly restricts ubiquitin signaling by an E2 suggests that noncatalytic E2 inhibition may be a more broadly conserved function of otubains than has been previously appreciated.

Our phylogenetic and proteomic DUB analyses support previous observations that microsporidia have undergone dramatic evolutionary reduction relative to other eukaryotes^57,58^. It will be important to understand whether a reduction in encoded DUBs reflects: i) reduced reliance on ubiquitin signaling for cellular function; ii) a relaxation of the substrate specificities of existing microsporidial DUBs to offset the reduced DUB complement; or iii) a bias toward degradative ubiquitination, which would presumably render dynamic chain editing less critical. Although microsporidia are most closely related to Fungi, many Fungi, including *S. cerevisiae*, lack otubain members^59^. Interestingly, otubain DUBs can be readily identified in other *Encephalitozoon* species infecting humans as well as in some closely related genera, whereas they are lost in other, more distant human pathogenic microsporidia, such as *Enterocytozoon bieneusi, Anncaliia algerae, Trachipleistophora hominis* (unpublished observations). The significance of this to host or tissue tropism, host cell infection, or parasite replication is not yet clear, but could potentially yield key insights into functional specialization among human microsporidial parasites.

The isolation of Ubc1/UBE2K as the sole E2 in coprecipitation experiments in yeast, as well as the lack of affinity for or inhibition of other human E2 enzymes (e.g., UBE2D1, Fig. 3C; UBE2N−UBE2V1, Supplementary Fig. S4) suggests that ehOTUB1 may have evolved exquisite selectivity for UBE2K, although this has not yet been comprehensively tested. Interestingly, although all E2 enzymes harbor a conserved UBC fold, UBE2K uniquely possesses a polyalanine loop (Supplementary Fig. S8) that we have demonstrated via mutagenesis makes intimate contact with ehOTUB1. In contrast, other human E2s contain comparatively bulky residues in this loop. It is thus appealing to speculate that ehOTUB1 may discriminate UBE2K based on the size of the side chains of the amino acids comprising this loop. It is likely that this is not the sole discriminatory feature, however, because yeast Ubc1 contains comparatively bulkier proline and valine at these positions, and ehOTUB1 retains interaction with Ubc1 (Fig. 3B,C). Structural studies will be invaluable to refine the interaction model presented here and to better understand how ehOTUB1’s E2 preference is enforced.

Although our modeling, mutagenesis, and functional assays suggest that ehOTUB1 and UBE2K interact using the same surfaces as observed for human OTUB1 and UBE2N^44,45^, ehOTUB1 employs a starkly different mechanism of inhibition (Fig. 8). Human OTUB1 preferentially binds the thioesterified form of UBE2N (Ub∼UBE2N, Fig. 8B) by virtue of interactions between its conserved N-terminal helix and the thioesterified ubiquitin. Further, this inhibition is potentiated by docking of a second ubiquitin in the human OTUB1 active site. In contrast, ehOTUB1 appears to favor the unmodified form of UBE2K (Fig. 8C), and inhibition of UBE2K is not obviously potentiated by docking of ubiquitin into the ehOTUB1 active site. Thus, ehOTUB1 acts earlier in the ubiquitin conjugation cascade, interfering with interaction between E1 and E2 to prevent E2 charging with ubiquitin.

**Figure 8:**
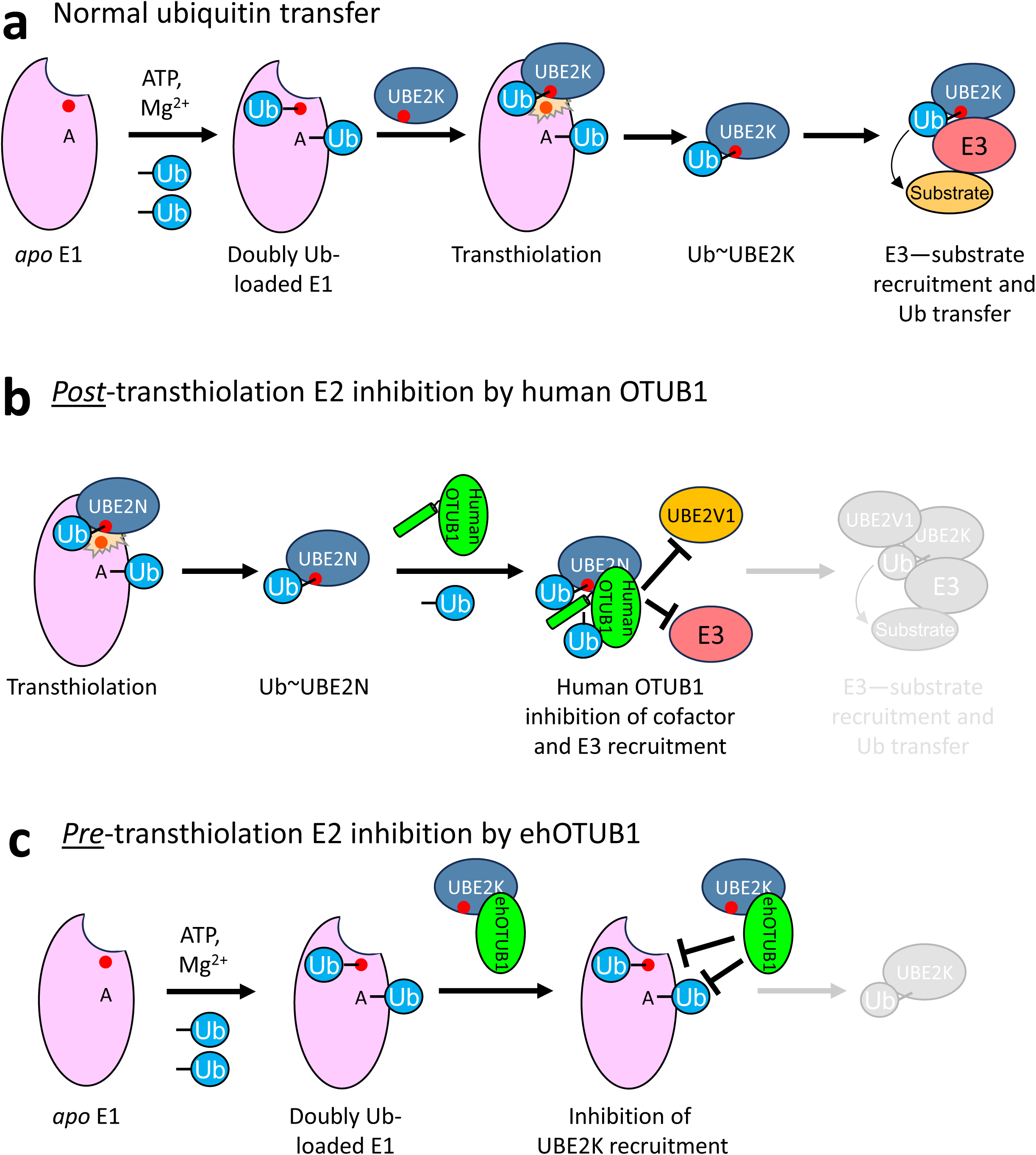
Comparison of pre- and post-transthiolation E2 inhibition mechanisms. ***A***, Cartoon of E2 transthiolation and subsequent E3-mediated ubiquitin transfer to substrates. ***B***, Human OTUB1 mediates E2 inhibition at post-transthiolation steps due to its preference for thioesterified E2, conferred by its N-terminal ubiquitin-binding helix. This configuration blocks recruitment of cofactor UBE2V1 and E3 ligases. ***C***, Pre-transthiolation inhibition of UBE2K by ehOTUB1. Steric conflict mediated in part between ehOTUB1 and ubiquitin bound in the E1 adenylation site prevents transthiolation of UBE2K.

Microsporidial genomes have undergone extensive evolutionary compaction; as a result, many proteins are significantly smaller than their counterparts in free-living eukaryotes. The distinct mechanism of E2 inhibition by ehOTUB1 that is independent of a ubiquitin-binding helix is a fascinating example of how microsporidial proteins often dispose of structural elements without compromising biological function. Whether this pre-transthiolation mechanism results in more robust inhibition of E2 activity than the post-transthiolation mechanism utilized by human OTUB1 is not yet clear. However, this work suggests that microsporidia may have evolved additional novel mechanisms to modify ubiquitin signaling with relevance to parasite physiology or host cell manipulation that warrant further exploration.

Although the biological function(s) of UBE2K remain rather poorly understood, multiple studies have shown that UBE2K catalyzes ubiquitination and degradation of cyclin B^60–62^, and it is well-established that cyclin B degradation drives cell cycle progression. Interestingly, infection by species of the *Encephalitozoon* genus, including *E. hellem*, causes cell cycle arrest accompanied by accumulation of cyclin B^63,64^. Upregulation of cyclin B has also been described upon infection of insect cells with species of the closely related genera *Nosema* and *Vairimorpha*^64^, whose genomes also encode ehOTUB1 orthologs. Notably, members of all three genera typically infect cells of the host gastrointestinal tract, and the epithelial cells lining the gastrointestinal tract constantly self-renew via cell division^65^. Thus, ehOTUB1 and its orthologs may function during infection to slow the replacement of these epithelial cells enough for sufficient parasite replication prior to the cells sloughing from the gastrointestinal surface via stabilization of cyclin B. Future studies aim to address this hypothesis.

## Experimental Procedures

### Cloning and plasmids

Plasmids used in this study are listed in Table S1. Plasmids were constructed using standard molecular cloning approaches with TOP10 F’ (Thermo Fisher Scientific) as the host. The ehOTUB1 coding sequence was cloned from *Encephalitozoon hellem* chromosomal DNA by PCR. Point mutations were introduced via site-directed mutagenesis. All coding sequences were verified by Sanger sequencing prior to use. Complete plasmid sequences and construction details are available upon request.

### Phylogenetic analysis of putative *E. hellem* deubiquitinating enzymes

Putative DUBs encoded by *E. hellem* were identified using the Protein BLAST suite from the National Center for Biotechnology Information (https://blast.ncbi.nlm.nih.gov/Blast.cgi?PAGE=Proteins) using the catalytic domains of members of each of the eight recognized DUB families (JAMM, MINDY, MJD, OTU, UCH, USP, VTD, and ZUFSP) as queries against the non-redundant protein sequence database, limited to *Encephalitozoon hellem* (taxid: 27973). The three-dimensional structures of the catalytic domains of hits were modeled using AlphaFold 3 (https://alphafolderserver.com), and the model with the highest confidence was inspected for conservation and reasonable three dimensional arrangement of catalytic residues. Assignment as orthologs of known *S. cerevisiae* DUBs was conducted by systematic pairwise sequence comparison using MAFFT (https://mafft.cbrc.jp/alignment/server/).

### Human cell culture, transient transfection, and immunoblotting

293T (ATCC) and Flp-In T-REx 293 cells (Thermo-Fisher) were maintained in DMEM supplemented with 10% FBS and 100 U/mL penicillin-streptomycin, and 1% (v/v) *L*-glutamine in a humidified incubator at 37°C with 5% CO_2_. Flp-In T-REx 293 cells additionally received 5 µg/mL blasticidin S and 100 µg/mL Zeocin (Thermo-Fisher). Cells were transiently transfected using jetPRIME reagent, followed by addition or not of 1 μg/mL doxycycline HCl to induce expression of the target protein. For measurement of the effects of ehOTUB1 on polyubiquitin in human cells, after induction of ehOTUB1 expression for 24 hours as above, 20 μM MG132 or DMSO vehicle was added for four hours prior to harvesting.

For immunoblotting analysis, transfected cells were harvested after 24 hours of expression by scraping into ice-cold PBS, centrifugation to pellet cells, and resuspension of the cell pellet in RIPA buffer (50 mM Tris-Cl, pH 8.0, 150 mM NaCl, 2 mM EDTA, 10 mM NaF, 1% v/v NP-40, 0.1% w/v SDS) supplemented with 1 mM AEBSF,and 20 mM *N*-ethylmaleimide to inactivate deubiquitinating enzymes. After incubation on ice with frequent vortexing for 30 minutes, the extracts were clarified by centrifugation at 21,000 x *g*, 4°C for 10 minutes. Equal amounts of protein were then separated by SDS-PAGE, transferred to PVDF membranes for one hour at 100V, 4°C, and subjected to immunoblotting with antibodies against K48-linked ubiquitin (Cell Signaling #4289; 1:2000), anti-FLAG M2 (Wako #012-22384; 1:2000), or anti-β-actin (Sigma #A5441, 1:10,000) and horseradish peroxidase-conjugated secondary antibodies (Cytiva) using enhanced chemiluminescence reagent^66^ and imaged on a Bio-Rad Chemidoc MP.

### Shotgun proteomics analysis of *E. hellem* deubiquitinating enzymes and protein identification

Uninfected 293T cells or 293T cells infected with *E. hellem* (∼80% infected at time of harvest) were harvested from an 80% confluent T182 flask, washed twice with ice-cold phosphate-buffered saline (PBS) and pelleted by centrifugation at 1000 x *g*, 4°C. Cells were lysed on ice in a modified RIPA buffer (50 mM Tris-HCl pH 8.0, 150 mM NaCl, 10 mM sodium fluoride, and 2 mM EDTA) supplemented with protease and phosphatase inhibitor cocktails. Following resuspension, 1% v/v NP-40 and 0.1% w/v SDS were added to facilitate membrane solubilization and further release of cellular contents. Lysis proceeded for 30 minutes on ice with intermittent vortexing. Lysates were clarified by centrifugation at 20,000 × *g* for 10 minutes at 4°C. The supernatant was collected, protein concentration was determined using the Bradford assay, and lysates were either stored at −80°C or used immediately for proteomics.

For pulldown of deubiquitinating enzymes, 100 μL of Ub-prg–bound HaloLink resin was pre-washed three times with 500 μL of Ub/DUB buffer (50 mM Tris-HCl, pH 7.5, 150 mM NaCl). Then, 10 mg of cell lysate was incubated with 100 μL of prepared resin for 1 hour at 37°C with gentle agitation. Following incubation, the resin was pelleted by centrifugation (1,500 × *g*, 4°C, 60 seconds), and the unbound material was collected. The resin was then washed with 500 μL of buffer 1 (350 mM NaCl, 50 mM Tris-HCl, pH 7.5), followed by 500 µL of buffer 2 (150 mM NaCl, 50 mM Tris-HCl, pH 7.5). For elution, the resin was incubated in 100 μL of Ub/DUB buffer containing 10 μg of purified HRV3C protease for 2 hours at 4°C with nutation to release trapped Ub-DUB conjugates. Following digestion, the sample was centrifuged at 1,500 × *g* for 1 minute, and the supernatant was collected. To remove the 3C protease, the eluate was incubated with 20 μL of Ni-NTA resin for 30 minutes at 4°C. The mixture was then loaded onto a microcentrifuge spin filter and centrifuged at 1,500 × *g* to separate the flowthrough from the resin. The final eluate was collected for downstream analysis.

Eluates were mixed 1:1 (v/v) with 10% SDS, 100 mM tetraethylammonium bromate (TEAB), pH 8.5 to a final volume of 23 µL containing approximately 100 µg of protein. Cysteine reduction and alkylation was performed via 55°C incubation with 5 mM TCEP for 15 minutes, followed by addition of 20 mM methylmethanethiosulfonate and incubation at room temperature for 10 minutes. The sample was then acidified by addition of 2.5 µL of phosphoric acid (∼2.5% w/v final concentration), followed by the addition of 165 µL of S-Trap binding/wash buffer (90% methanol, 100 mM TEAB, pH 7.5). The acidified sample was loaded onto an S-Trap micro column (ProtiFi) and centrifuged at 4,000 × *g* for 30 seconds to bind proteins. The column was washed three times with 150 µL binding/wash buffer, followed by a final centrifugation at 4000 x *g* for one minute to remove residual wash buffer. Twenty microliters of sequencing-grade trypsin (1:10 w/w enzyme : protein ratio) in 50 mM TEAB, pH 7.1 was added to the column, and incubated at 47 °C for 2 hours. Peptides were eluted sequentially with 40 µL of 50 mM TEAB pH 8.5, 0.2% (v/v) formic acid in water, and 50% (v/v) acetonitrile in water, with eluates collected by centrifugation at 4,000 × *g* for 1 minute after each step. The eluates were pooled, dried using a SpeedVac, and resuspended in 20 µL of 0.1% (v/v) formic acid for analysis.

Mass spectrometry was performed at the Translational Science Laboratory in the College of Medicine at Florida State University. An externally calibrated Thermo Scientific Orbitrap Exploris 480 mass spectrometer was used in conjunction with the Thermo Scientific EASY-nLC 1200 system. Trypsin-digested sample was loaded onto the trap column (Acclaim PepMap 100, 100 μm × 2 cm, nanoViper; Thermo Scientific). The flow rate was set to 300 nL/min for separation on the analytical column (Acclaim pepmap RSLC, 75 μM × 15 cm, nanoViper; Thermo Scientific). Mobile phase A was composed of 99.9% H2O containing 0.1% formic acid and mobile phase B was composed of 95% acetonitrile, 4.9% H2O, and 0.1% formic acid. A 60-minute gradient from 5% to 45% mobile phase B was performed. The LC eluent was directly nanosprayed into the Exploris 480 mass spectrometer. During chromatographic separation, the Exploris 480 was operated in a data-dependent mode under the direct control of the Thermo Excalibur 4.4.16.14 (Thermo Scientific). MS data were acquired using the following parameters: 30 data-dependent collisional-induced-dissociation (CID) MS/MS scans per full scan (350 to 1700 m/z) at 120,000 resolution in profile mode. MS2 were acquired in centroid mode at 15,000 resolution. Ions with a single charge, more than 7 charges, or unassigned charges were excluded. A 15-second dynamic exclusion window was used. All measurements were performed at room temperature, and three technical replicates were run for each sample. The raw files were analyzed using Thermo Proteome Discoverer (version 2.5.0.400) software package with SequestHT search node using the *S. cerevisiae* reviewed (Swiss-Prot) 2024-09-05 fasta database downloaded from uniprot.org with a custom-added sequence for 3xFLAG-tagged ehOTUB1 for yeast experiments, or using the human reviewed fasta file (uniprot.org ID 9606) and *E. hellem* 2024-09-05 FASTA database (uniprot.org).] and the Percolator peptide validator. The resulting .msf files were further validated by the proteome validator software Scaffold v5.3.0 (Portland, OR, USA).

### Expression and purification of ehOTUB1 and mutants

WT ehOTUB1 and all mutants were expressed as a C-terminal fusion to 12His-*Brachypodium distachyon* SUMO (12His-bdSUMO) in LOBSTR(DE3) cells harboring pRARE2. Cells were resuspended in Tris-NTA buffer (50 mM Tris-Cl, pH 7.5, 500 mM NaCl, 0.2% Triton X-100, 20 mM imidazole, 10% glycerol, 5 mM β-mercaptoethanol), lysed by two passes through an Avestin Emulsiflex-C5 high-pressure homogenizer, and the resultant lysates were clarified by centrifugation at 30,000 x *g* for 20 minutes at 4°C. Clarified lysates were bound to Ni-NTA resin for 30 minutes at 4°C, washed extensively with Tris-NTA wash buffer (50 mM Tris-Cl, pH 7.5, 500 mM NaCl, 0.2% Triton X-100, 70 mM imidazole, 10% glycerol, 5 mM β-mercaptoethanol), and subjected to on-column cleavage in Tris-NTA wash buffer supplemented with 2 mM TCEP and 500 nM *Brachypodium distachyon* SENP1 for two hours at 4°C. The cleaved protein was collected, concentrated using a 10 kDa MWCO filter (Amicon), and further purified via gel filtration (Superdex 75 pg or a hand-packed 120 mL Superose 6 pg column) in Ub/DUB buffer (50 mM Tris-Cl, pH 7.5, 100 mM NaCl, 0.1 mM TCEP). Proteins used for crosslinking experiments were instead separated in Ub/DUB buffer lacking reductant. Peak fractions were pooled, concentrated, and snap-frozen in liquid nitrogen for storage at −80°C.

### Purification of ubiquitin and mutants

Ubiquitin and all mutants were expressed via autoinduction in LOBSTR(DE3) cells harboring pRARE2. Cell pellets were resuspended in Ub-CatA buffer (50 mM ammonium acetate, pH 4.5) supplemented with 1 mM PMSF, and lysed via homogenization as above. The lysate was then heated to 80°C for 20 minutes to precipitate most *E. coli* proteins. The precipitated lysate was cooled in an ice-water bath and centrifuged at 30,000 x *g* for 20 minutes, 4°C. The supernatant was applied to a 20 mL HiLoad SP cation exchange column equilibrated in Ub-CatA buffer and eluted with a linear gradient of Ub-CatB buffer (50 mM ammonium acetate, pH 4.5, 1 M NaCl). Peak fractions were pooled, dialyzed overnight into 50 mM Tris-Cl, pH 7.5, concentrated to 3-6 mM in a 3 kDa MWCO filter (Amicon), and stored in single-use aliquots at −80°C.

Ubiquitin, NEDD8, SUMO-1, SUMO-3, and ISG15 residues 79-155 (ISG15^79–155^) were expressed as N-terminal intein-chitin binding domain fusions as done previously^67^. Cells were lysed as above in Chitin lysis buffer (50 mM HEPES-acetate, pH 6.5, 50 mM sodium acetate, 500 mM NaCl, 0.2% Triton X-100), and lysates were clarified by centrifugation at 30,000 x *g*, 20 minutes, 4°C. The supernatant was applied to chitin resin (NEB) for 2 hours at 4°C, washed extensively in Chitin lysis buffer, and the resin was then equilibrated in chitin cleavage buffer (50 mM HEPES-acetate, pH 6.5, 50 mM sodium acetate, 75 mM NaCl, 100 mM 2-mercaptoethanesulfonate sodium). The resin was then incubated overnight at 37°C to release the ubiquitin/ubiquitin-like protein as a C-terminal thioester.

For synthesis of Ub-AMC, the thioester was then concentrated to approximately 3 mg/mL and the procedure of Wilkinson *et al.*^67^ was followed exactly as described. The Ub-AMC conjugate was purified by cation exchange and dialyzed as described above for ubiquitin. The Ub-AMC eluted at a higher NaCl concentration than the Ub_1-75_ thioester hydrolysis product. Ub-AMC was concentrated to 500 μM, and stored in small aliquots at −80°C. To produce C-terminal propargylamide forms of each Ubl, the thioester form was adjusted to pH 8.0 with 1 M NaOH, and propargylamine (Sigma) was immediately added to 250 mM. After incubation for four hours at 25°C, the protein-propargylamide was separated from unreacted propargylamine by gel filtration on a Sephacryl S-200 column equilibrated in Ub/DUB buffer. Peak fractions were pooled, concentrated to approximately 200 μM, and stored at −80°C until use.

### Preparation of native ubiquitin chains

M1-linked ubiquitin chains were expressed as C-terminal 12His-bdSUMO fusions and purified as described above for ehOTUB1. K27-linked Ub_2_ chains were purchased from Life Sensors. All other ubiquitin chains were synthesized using linkage-specific E2 or E3 enzymes^68^. The enzymes and conditions are listed in Table S2. After synthesis, chain assembly reactions were diluted 10-fold in Ub-CatA buffer, applied to a 20 mL HiLoad SP column, and eluted with a linear gradient of Ub-CatB buffer. Fractions containing Ub_2_, Ub_3_, or Ub_4_ chains were identified by SDS-PAGE, pooled, dialyzed overnight into 50 mM Tris-Cl, pH 7.5, concentrated to approximately 500 μM in a 3 kDa MWCO filter (Amicon), and stored at −80°C.

### Preparation of proximal-mutant K48-linked diubiquitin chains

K48-linked Ub_2_ chains harboring mutations in the proximal unit were synthesized using a 1:1 ratio of Ub(K48R) and either Ub(D77) or Ub(D77) harboring additional mutations. The K48R and D77 mutations ensure that Ub(K48R) and Ub(D77) occupy the distal and proximal positions within the diubiquitin chain, respectively. Conditions for synthesis, purification, and storage were otherwise identical to those for native ^K48^Ub_2_ (Table S2).

### Synthesis of branched triubiquitin

Branched Ub_3_ chains in which two distal ubiquitin moieties are linked to a single proximal ubiquitin were synthesized using Ub(D77) as the proximal ubiquitin, and with Ub(K11R,K48R), Ub(K11R,K63R), or Ub(K48R,K63R) as the distal ubiquitins to form branched [Ub_2_]–^11,48^Ub, [Ub_2_]–^11,63^Ub, and [Ub_2_]–^48,63^Ub, respectively (Table S2). Chains were then purified, dialyzed, concentrated to approximately 200 μM, and stored as described above. Linkages were verified using recombinant Cezanne, human OTUB1, and AMSH*^69^, which preferentially hydrolyze K11, K48, and K63 linkages, respectively.

### Deubiquitination assays

A 2x solution of WT or the indicated ehOTUB1 mutants were preincubated in Ub/DUB buffer supplemented with 5 mM DTT for 5 minutes at room temperature before mixing with an equal volume of 2x ubiquitin chains to initiate proteolysis (final concentrations: 500 nM ehOTUB1; 10-20 μM ubiquitin chains). Aliquots were collected at the indicated timepoints and mixed with an equal volume of 2x Laemmli loading buffer to terminate the reaction, and stored at −20°C until analysis.

### Ubiquitin conjugation assays

Ubiquitin conjugation reactions contained 1 μM E1, 5 μM human UBE2K or 1 μM human Ubc13:Mms2, and a mixture of 1 part Oregon Green 488-labeled ubiquitin and nine parts unlabeled ubiquitin (total ubiquitin concentration was 300 μM) in 50 mM Tris-Cl, pH 8.0, 10 mM MgCl_2_, 1 mM DTT. Buffer, ehOTUB1, or the ehOTUB1(C51A) were then added and incubated for 5 minutes prior to addition of 10 mM ATP to initiate the reaction. After incubation at 37°C for 60 minutes, the reaction was terminated by addition of an equal volume of 2x Laemmli loading buffer, and the samples were stored at −20°C until analysis.

### Size exclusion interaction analyses

For analysis of interaction between ehOTUB1, UBE2K, and their respective mutants, the proteins or their mixtures were incubated at 4°C for two hours before separation on a Superdex 75 pg column equilibrated in Ub/DUB buffer + 0.1 mM TCEP. For K48-Ub_2_ binding to ehOTUB1 and mutants, K48-Ub_2_ was incubated with a 1.2-fold molar excess of ehOTUB1(C51A) or mutants for one hour at room temperature, followed by separation on a Superose 6 Increase 10/30 column equilibrated in Ub/DUB buffer + 0.1 mM TCEP.

### UBE2K thioesterification assays

Purified UBE2K (final assay concentration 800 nM) was pre-incubated with 5 µM ehOTUB1(C51A), ehOTUB1(C51A,L94W,F99W) or buffer control for 15 minutes at room temperature before mixing with an equal volume of E1 (40 nM final), ubiquitin (25 µM), and ATP (2 mM). Zero-minute control was prepared identically, but with E1 mixture lacking ATP. At the indicated times after mixing, 20 µL of the reaction was added to 10 µL of 5x non-reducing Laemmli buffer, mixed, and kept on ice until completion of the assay. Where indicated, DTT was added to 33 mM, and samples were then heated at 42°C for three minutes before separation on 13% Bis-Tris gels and Coomassie staining.

### E1-E2 crosslinking assays

The sole cysteine (catalytic Cys92) in UBE2K was first activated with 1.25 mM 2,2’-dipyridyldisulfide for 30 minutes at room temperature, followed by desalting into Ub/DUB buffer to remove unreacted 2,2’-dipyridyldisulfide. Single-use aliquots were made and snap-frozen at −80°C until use. E1 was reduced on ice with 1 mM TCEP for 30 minutes before desalting to remove excess TCEP and snap-frozen as above. For crosslinking assays, 10 µL of 2 µM E1 was mixed with 10 µL of 5 µM activated UBE2K for 60 seconds at room temperature (final concentrations 1 and 2.5 µM, respectively) before quenching with 10 µL of 5x non-reducing Laemmli buffer and immediate analysis by non-reducing SDS-PAGE. In experiments also containing ehOTUB1 (final concentration 10 µM) or ubiquitin (20 µM), ehOTUB1 was pre-incubated with UBE2K whereas ubiquitin was pre-incubated with E1 for 10 minutes prior to assay as above. Where indicated, 1 µL of 1 M dithiothreitol was added to the reaction after crosslinking to reduce the E1-UBE2K disulfide bond before SDS-PAGE.

### Pseudoatomic modeling of ehOTUB1 alone and in complex with other proteins

Structure predictions of ehOTUB1 or ehOTUB1 in complex with human UBE2K were run using the AlphaFold 3 server (http://www.alphafoldserver.com). Agreement between the top five models was assessed using the rms function in Pymol (http://www.pymol.org). Modeling of the ehOTUB1−UBE2K AlphaFold structure onto published structures was performed using either the Matchmaker command in ChimeraX (https://www.cgl.ucsf.edu/chimerax/) or the Align command in Pymol with the UBE2K chain from the AlphaFold model and the relevant E2 enzyme in the published structure.

## Acknowledgements

The authors thank Jonathan Snow (Barnard College) for sharing *E. hellem* and for helpful discussions. We thank members of the Tomko, Yanchang Wang, and Hong-Guo Yu labs for constructive feedback on the work. We are indebted to MicrosporidiaDB (http://www.microsporidiadb.org) for curating information about *E. hellem* and other microsporidia. This work was supported by R01 GM116800 and an FSU Council for Research and Creativity Seed Grant to R.T.

## Author contributions

R.T. conceived the studies with input from T.L. and A.N.; T. L., performed DUB proteomics experiments; C.V. performed mass spectrometry and associated data analysis; N.C. and T.B. performed yeast genetic experiments; A.C. performed human cell experiments; T.L., L.C., and R.T. purified all proteins and performed biochemical experiments. R.T. and T.L. wrote the first manuscript draft; all authors reviewed the manuscript. R.T. acquired funding and oversaw the project.

## Figure Legends

**Supplemental Figure S1:**
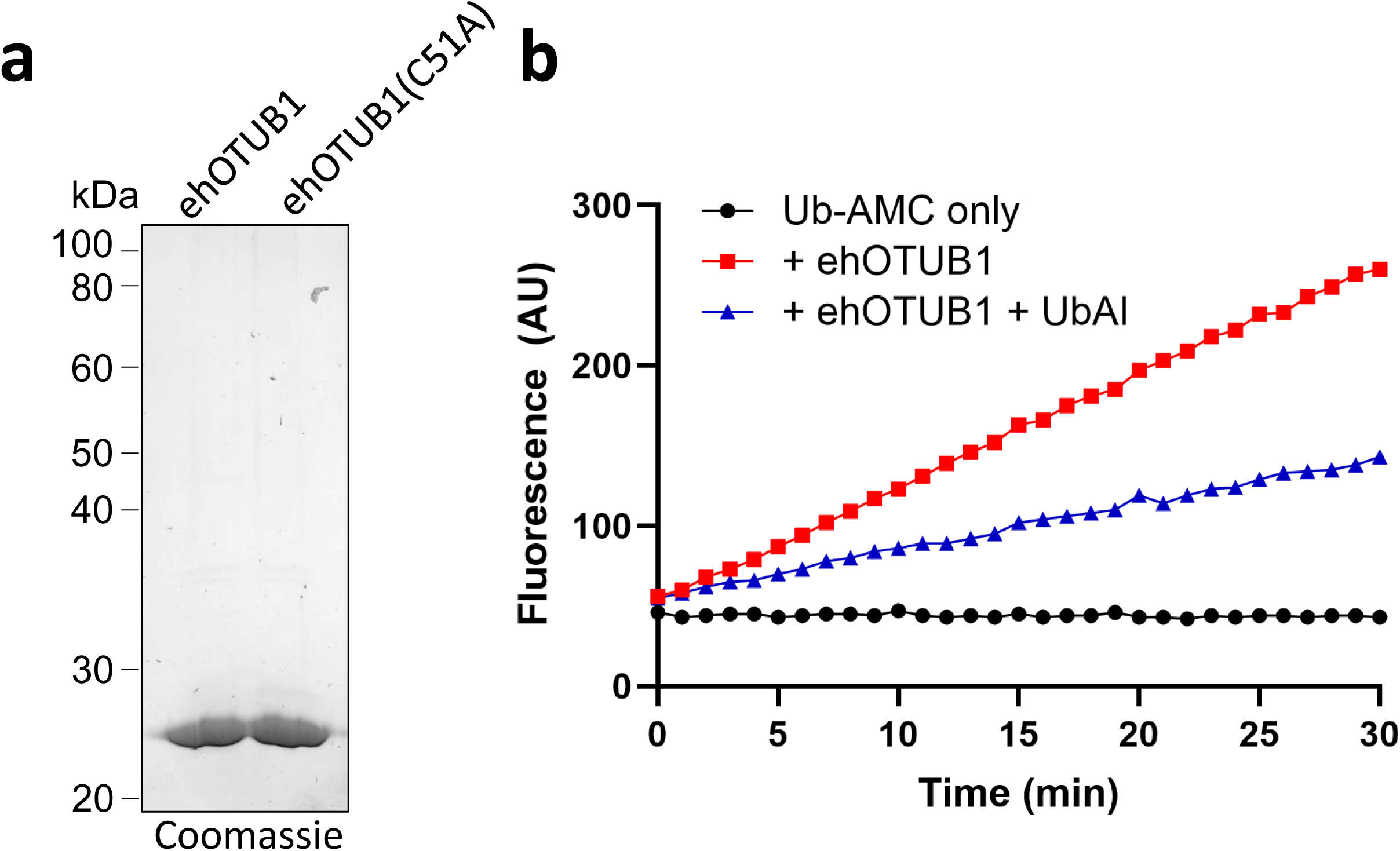
Purification of ehOTUB1 and assessment of ubiquitin hydrolase activity. ***A***, SDS-PAGE analysis of purified recombinant ehOTUB1 and ehOTUB1(C51A). ***B***, Ubiquitin aldehyde inhibits ehOTUB1-mediated Ub-AMC hydrolysis. Ub-AMC hydrolysis by ehOTUB1 was measured in the presence or absence of competitive inhibitor ubiquitin aldehyde.

**Supplemental Figure S2:**
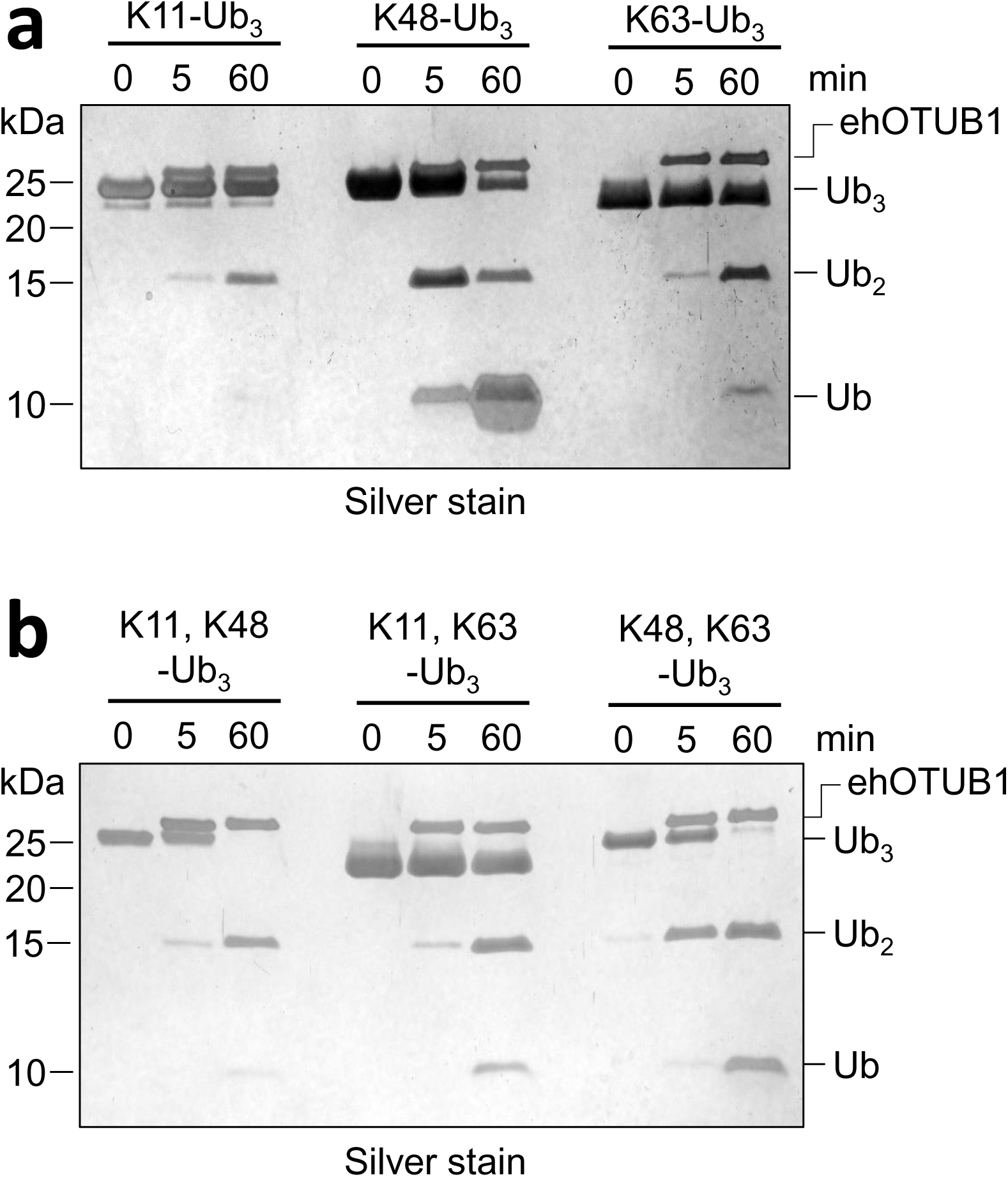
Processing of more complex ubiquitin chains by ehOTUB1. ***A***, K11-, K48-, or K63-linked triubiquitin were incubated with 500 nM ehOTUB1 for the indicated times before analysis by SDS-PAGE. No improvement of K11- or K63 linkage activity was observed with Ub_3_. Similar results were obtained with Ub_4_ (not shown). ***B***, ehOTUB1 cleaves branched ubiquitin chains. Triubiquitin chains in which the proximal ubiquitin bears two ubiquitin linkages via the indicated lysines were incubated with 500 nM ehOTUB1 for the times shown before analysis by SDS-PAGE.

**Supplemental Figure S3:**
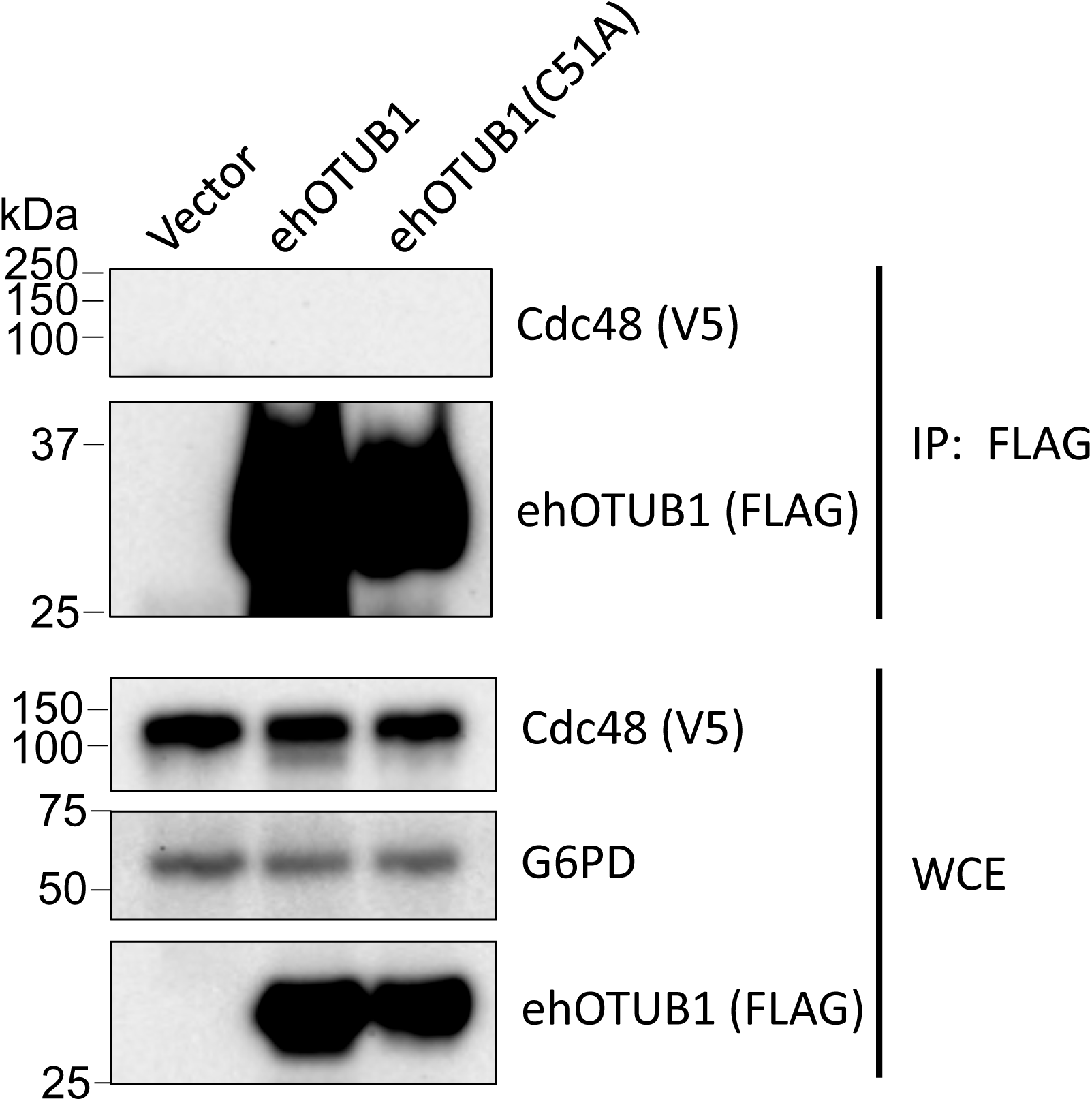
ehOTUB1 does not interact detectably with Cdc48. A yeast strain in which Cdc48 is expressed from its native chromosomal locus with a C-terminal V5 tag was transformed with plasmids encoding the indicated forms of 3xFLAG-ehOTUB1, and extracts were subjected to FLAG immune precipitation. The FLAG immunoprecipitates and whole cell extracts were then probed for V5 (Cdc48) or FLAG. G6PD is shown as a loading control.

**Supplemental Figure S4:**
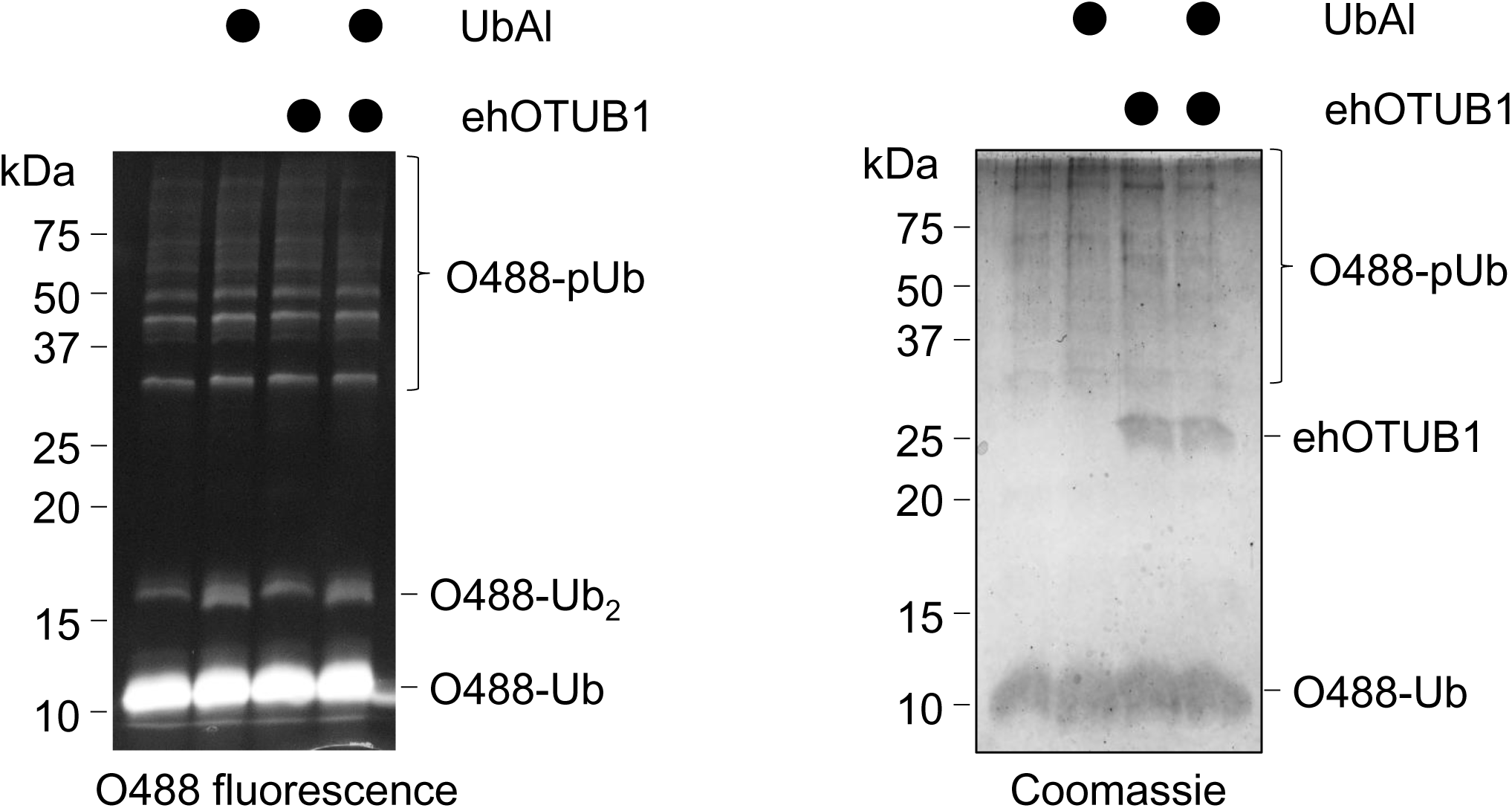
ehOTUB1 does not inhibit UBE2N−UBE2V1 ubiquitin conjugating activity. ehOTUB1 (500 nM) was pretreated or not with 500 nM ubiquitin aldehyde, and then added to 100 nM E1, 400 nM UBE2N−UBE2V1, 5 µM Oregon Green 488-Ub, and 0.6 mM DTT. The reaction was initiated with 10 mM ATP, and after 3 hours the reaction was terminated by the addition of loading buffer. Ubiquitin conjugates were then visualized via Oregon Green 488 fluorescence, and total proteins by Coomassie staining.

**Supplemental Figure S5:**
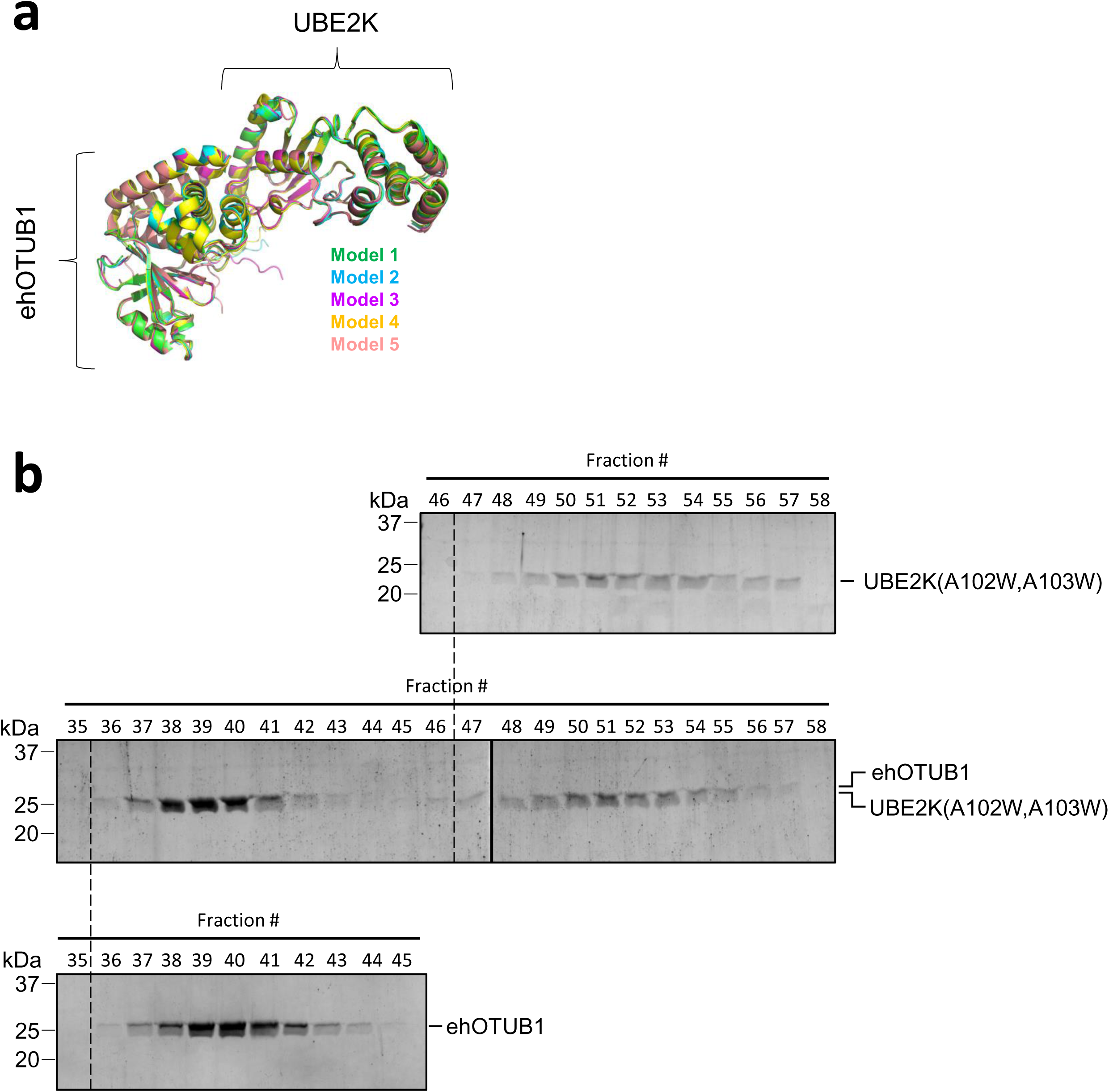
Interaction between ehOTUB1 and UBE2K predicted by Alphafold modeling. ***A***, Superimposition of the top five AlphaFold models of the ehOTUB1−UBE2K complex. RMSD was < 0.5 Å between each pair of models. ***B***, SDS-PAGE analysis of size exclusion fractions from the experiment shown in Main Text Fig. 5D showing slowed migration of full-length UBE2K(A102W,A103W), and a lack of migration shift in ehOTUB1 or UBE2K(A102W,A103W) when the proteins were mixed prior to size exclusion chromatography.

**Supplemental Figure S6:**
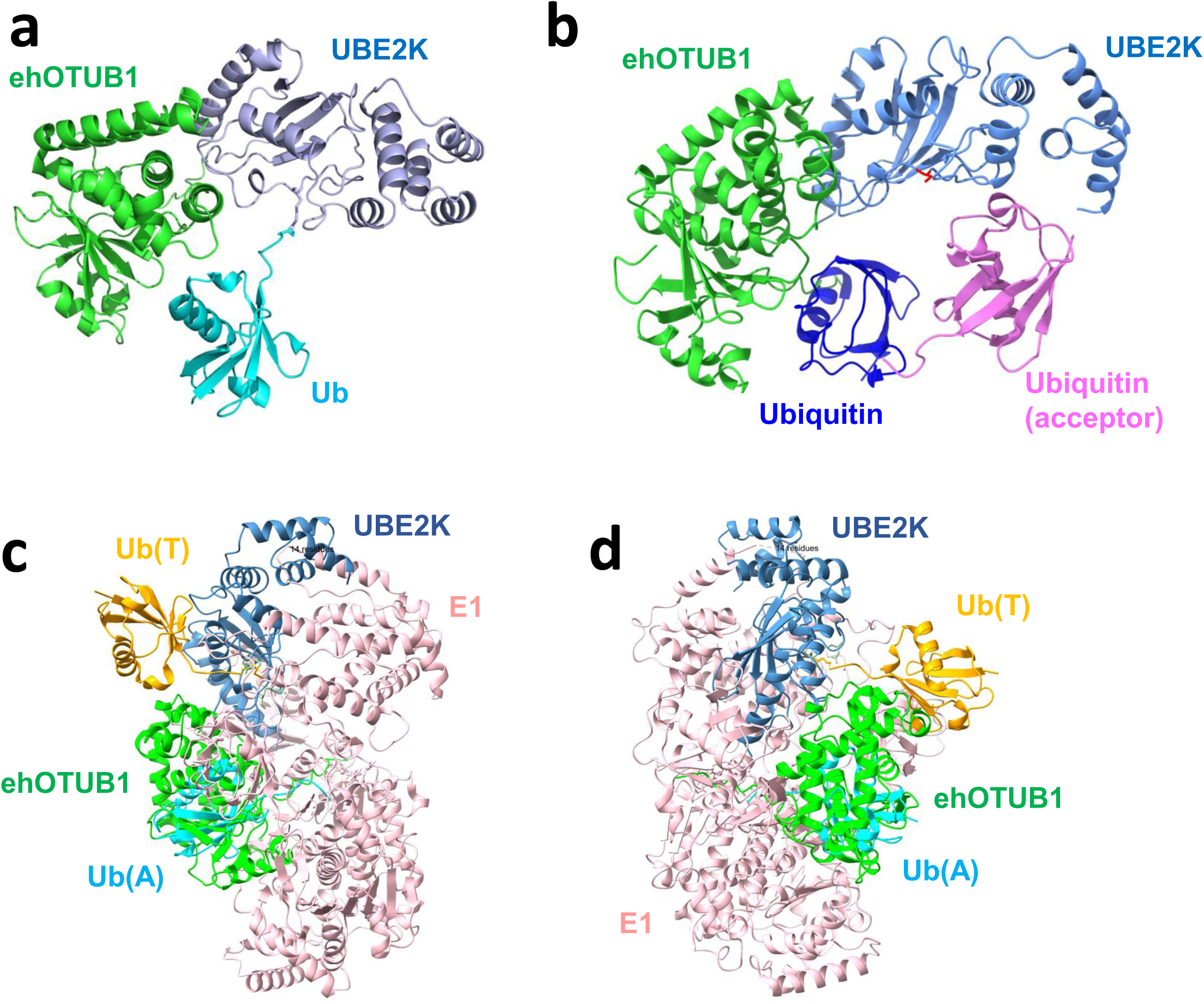
Binding of ehOTUB1 to UBE2K is not predicted to generate clash with thioesterified ubiquitin, with acceptor ubiquitin, or with the thioesterified ubiquitin bound to the E1 catalytic cysteine. *A*, ehOTUB1 can be superimposed onto the structure of ubiquitin thioesterified UBE2K (PDB 5DFL) without obvious steric conflict between ubiquitin (cyan) and ehOTUB1 (green). RMSD for UBE2K (cornflower blue) superimposition was 0.283 Å. *B*, ehOTUB1 is unlikely to directly interfere with acceptor ubiquitin recruitment. The ehOTUB1−UBE2K model colored as in (*A*) was superimposed onto a structure of UBE2K bound to acceptor K48-Ub_2_ (PDB 7OJX). No appreciable steric conflict was observed between K48-Ub_2_ (cyan and deep blue) and ehOTUB1 (green). *C* and *D*, ehOTUB1 does not conflict with thioesterified ubiquitin in E1-E2 *trans*thiolation intermediate structures. The ehOTUB1−UBE2K model was superimposed onto PDB structures 9B5D (*C*) or 9B5E (*D*). Ubiquitin in the E1 thioester site is shown in gold, ubiquitin in the E1 adenylation site is in cyan, E1 is in rose, ehOTUB1 in green, and UBE2K in cornflower blue.

**Supplemental Figure S7:**
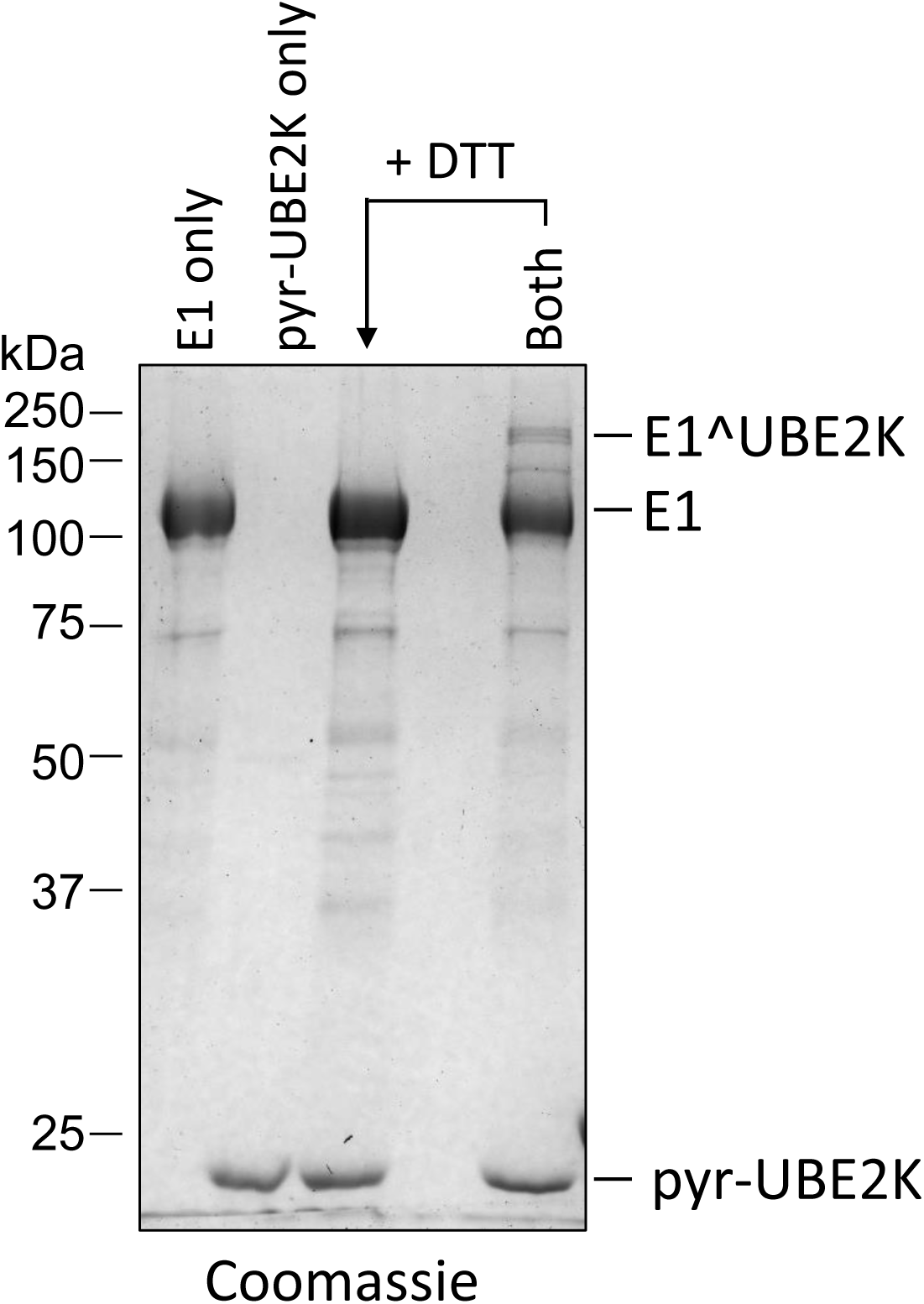
The E1-UBE2K crosslink depends on both E1 and pyr-UBE2K. E1, pyr-UBE2K, or the pair were incubated for 60 seconds before quenching with 5x non-reducing Laemmli buffer and analysis via non-reducing SDS-PAGE. Where indicated, 1 µL of 1 M DTT was added for 3 minutes prior to loading the sample.

**Supplemental Figure S8:**
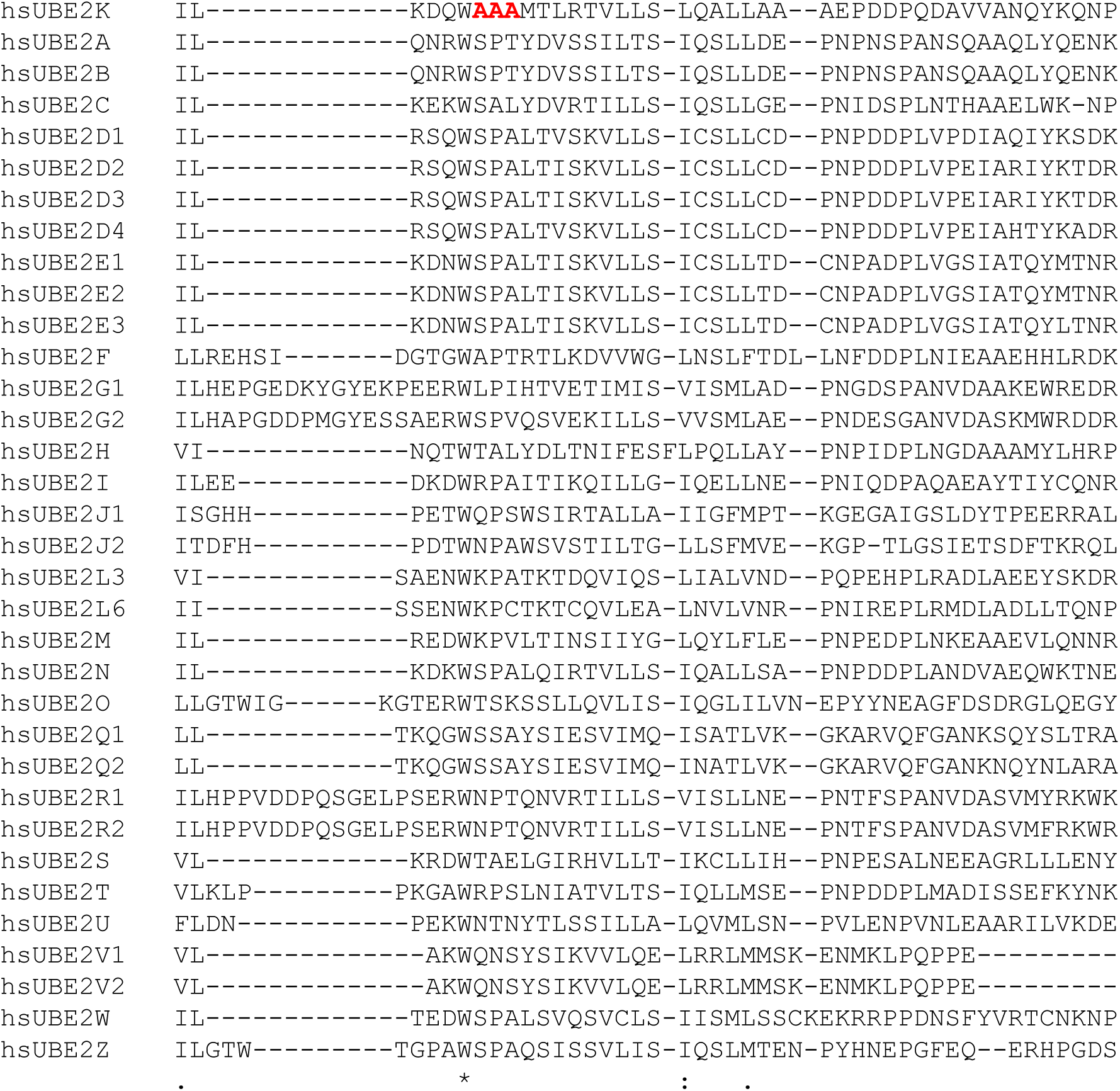
hsUBE2K uniquely contains a polyalanine stretch in an ehOTUB1-interacting loop. An alignment of all known human E2 enzymes to human UBE2K is shown, with the position of a polyalanine stretch in UBE2K shown in bold red font.

