## Supplementary material for "A microsporidial deubiquitinase blocks ubiquitin transfer from adenylated E1 to human UBE2K ubiquitin conjugating enzyme": Table 1

**Table 1: *E. hellem* deubiquitinating and ubiquitin-like protein deconjugating enzymes identified by phylogenetic search**

| DUB family | Putative *E. hellem* member | Apparent *S. cerevisiae* ortholog |
|---|---|---|
| JAMM | EHEL_110440 | *RPN11* |
| OTU | EHEL_050640 | None identified |
| USP | EHEL_030490 | *UBP15* |
| USP | EHEL_030570 | *UBP14* |
| USP | EHEL_051360 | *UBP8* |
| USP | EHEL_060860 | *UBP12* |
| USP | EHEL_070380 | *DOA4* |
| ULP (SUMO protease) | EHEL_050840 | *ULP1* |
| DeSI (putative SUMO protease) | EHEL_090660 | None identified |
