## Supplementary material for "A microsporidial deubiquitinase blocks ubiquitin transfer from adenylated E1 to human UBE2K ubiquitin conjugating enzyme": Table 2

**Table 2: *E. hellem* deubiquitinating enzymes identified from mass spectrometry of *E. hellem*-infected 293T cell extracts**

| **DUB family** | ***E. hellem* gene name** | **Apparent *S. cerevisiae* ortholog** | **Apparent human ortholog** |
|---|---|---|---|
| USP | EHEL_030490 | *UBP15* | *USP7* |
| USP | EHEL_030570 | *UBP14* | *USP5/Isopeptidase T* |
| USP | EHEL_070380 | *DOA4* | *USP8* |
| OTU | EHEL_050640 | None identified | None identified |
