## Supplementary Table S1 for "A microsporidial deubiquitinase blocks ubiquitin transfer from adenylated E1 to human UBE2K ubiquitin conjugating enzyme"

**Supplemental Table S1. Plasmids used in this study.**

| Plasmid | Genotype | Source |
|---|---|---|
| pRT45 | p424GPD | Mumberg et al, 1995 |
| pRT690 | pET28b-mmE1 | Addgene #32534 |
| pRT1219 | pcDNA5/FRT/TO | Thermo-Fisher Scientific |
| pRT1349 | pRSETA-hsUb | Gift from Judith Ronau |
| pRT1351 | pET26b-mmUb(K11R,K63R) | Gift from Judith Ronau |
| pRT1660 | pRT1254-hsUBE2K | This study |
| pRT1705 | pRT1254-hsUBE2SΔC | This study |
| pRT1723 | pET42b-hsUb(G76C)-6His | This study |
| pRT1738 | pETDuet-1-6His-3Cx-scUb(M1C) | This study |
| pRT1785 | pRT1254-scUbc1 | This study |
| pRT1880 | pOPINB-AMSH* | Addgene #66712 |
| pRT2281 | pRT2258-hsUBE2D1 | This study |
| pRT2323 | pTYB1-hsUb(1-75) | This study |
| pRT2399 | pTYB1-hsNEDD8(1-75) | This study |
| pRT2424 | pRT2258-hsUBE2N : hsUBE2V1 | This study |
| pRT2819 | pRT2258-ehOTU1 (EHEL_050640) | This study |
| pRT2821 | pOPINK-Cezanne(53-446) | Addgene #61581 |
| pRT2822 | pOPINB-OTUB1* | Addgene #65441 |
| pRT2823 | pOPINS-AREL1(436-823) | Addgene #66710 |
| pRT2824 | pOPINS-hsUBE3C(693-1083) | Addgene #66711 |
| pRT2825 | pMCSG17-NleL(170-782) | Addgene #66716 |
| pRT2826 | pRT2258-hsUBE2L3 | This study |
| pRT2853 | pRT2258-ehOTU1(C51A) | This study |
| pRT2858 | p424GPD-3xFLAG-ehOTU1 | This study |
| pRT2875 | p424GPD-*scOTU1* | This study |
| pRT2876 | p424GPD-*scOTU2* | This study |
| pRT2885 | p424GPD-3xFLAG-ehOTU1(C51A) | This study |
| pRT2937 | pRSETA-hsUb(K48R) | This study |
| pRT2938 | pRSETA-hsUb(D77) | This study |
| pRT2939 | pRSETA-hsUb(F4A,D77) | This study |
| pRT2940 | pRSETA-hsUb(L8A,D77) | This study |
| pRT2941 | pRSETA-hsUb(I36A,D77) | This study |
| pRT2946 | pGEX-4T-2-gp78(309-643) | A gift from Allan Weissman |
| pRT2957 | pRT2258-ehOTU1(9-222) | This study |
| pRT2959 | pRSETA-hsUb(I44A,D77) | This study |
| pRT2969 | pRT2258-ehOTU1(1-222) | This study |
| pRT2970 | pRT2258-ehOTU1(9-227) | This study |
| pRT2971 | pcDNA5/FRT/TO-3xFLAG-ehOTU1 | This study |
| pRT2972 | pcDNA5/FRT/TO-3xFLAG-ehOTU1(C51A) | This study |
| pRT2984 | pRT2258-ehOTU1(Y150A) | This study |
| pRT2985 | pRT2258-ehOTU1(D191A) | This study |
| pRT2986 | pRT2258-ehOTU1(K194A) | This study |

| | | |
|---|---|---|
| pRT2987 | pRT2258-ehOTU1(Y200A) | This study |
| pRT2993 | pTYB1-hsSUMO-1 | This study |
| pRT2994 | pTYB1-hsSUMO-3 | This study |
| pRT3007 | pTYB1-12His-Halotag-3Cx-hsUb(1-75) | This study |
| pRT3013 | pRT1998-hsUb(V70A,D77) | This study |
| pRT3022 | pTYB1-hsISG15(79-155) | This study |
| pRT3111 | pRT1998-hsUb(K48R,K63R) | This study |
| pRT3115 | pRT1998-hsUb(K48R,D77) | This study |
| pRT3128 | pRT1998-hsUb(K63R,D77) | This study |
| pRT3152 | pRT1998-hsUb(K11R,K48R) | This study |
| pRT3176 | pRT2258-ehOTU1(D92R) | This study |
| pRT3177 | pRT2258-ehOTU1(F99A) | This study |
| pRT3178 | pRT2258-ehOTU1(Y123A/D127R) | This study |
| pRT3179 | pRT2258-ehOTU1(Y168A) | This study |
| pRT3221 | pRT2258-ehOTU1(C51A,L94W,F99W) | This study |
| pRT3223 | pRT1254-hsUBE2K(A102W,A103W) | This study |
| pRT3316 | pRT2258-ehOTU1(L94W,F99W) | This study |
| pRT3323 | pRT2258-ehOTU1(9-222,C51A) | This study |
| pRT3324 | pRT2258-ehOTU1(1-222,C51A) | This study |
| pRT3325 | pRT2258-ehOTU1(9-227,C51A) | This study |
| pRT3326 | pRT2258-ehOTU1(C51A,Y150A) | This study |
| pRT3327 | pRT2258-ehOTU1(9-222,Y200A) | This study |
