## Supplementary Table S2 for "A microsporidial deubiquitinase blocks ubiquitin transfer from adenylated E1 to human UBE2K ubiquitin conjugating enzyme"

**Table S2. Conditions for preparation of linkage-specific ubiquitin chains**

| Linkage | [E1], µM | [E2], µM | [E3], µM | Ub, mM | [DUBs], µM | Temp (C), time |
|---|---|---|---|---|---|---|
| **K6-linked** | 1 | UBE2L3, 0.6 | NleL, 10 | 2.4 | OTUB1*, 2 | 37$^{o}$C o/n |
| **K11-linked** | 1 | UBE2SΔC, 10 | – | 2.4 | AMSH*, 1 | 37$^{o}$C o/n |
| **K29-linked** | 1 | UBE2L3, 10 | UBE3C, 30 | 2.4 | Cezanne, 0.4[1]<br>OTUB1*, 2 | 37$^{o}$C o/n |
| **K33-linked** | 1 | UBE2L3, 10 | AREL1, 20 | 2.4 | Cezanne, 0.4[1]<br>OTUB1*, 2<br>AMSH*, 2 | 37$^{o}$C o/n |
| **K48-linked** | 1 | UBE2K, 25 | – | 2.4 | – | 37$^{o}$C, 3 hrs |
| **K63-linked** | 1 | UBE2N-UBE2V1, 10 | – | 2.4 | – | 37$^{o}$C, 3 hrs |
| **[Ub$_2$] –$^{11,48}$Ub** | 1 | UBE2K, 20<br>UBE2SΔC, 20 | gp78, 20 | Ub(K11R,K48R), 1.6<br>Ub(D77), 0.8 | – | 37$^{o}$C o/n |
| **[Ub$_2$] –$^{11,63}$Ub** | 2 | UBE2SΔC, 20<br>UBE2N-UBE2V1, 20 | – | Ub(K11R,K63R), 1.6<br>Ub(D77), 0.8 | – | 37$^{o}$C o/n |
| **[Ub$_2$] –$^{48,63}$Ub** | 1 | UBE2K, 25<br>UBE2N-UBE2V1, 10 | – | Ub(K48R,K63R), 1.6<br>Ub(D77), 0.8 | – | 37$^{o}$C o/n |

All reactions were performed in 50 mM Tris-Cl, pH 8.0, 10 mM $MgCl_2$, 10 mM ATP, 0.6 mM DTT, and reactions were terminated by the addition of 10 mM DTT.

[1]Cezanne was added for one hour after quenching and prior to purification of chains.
